# Membrane Thickness Strain from Protein Inclusions: A Multiscale Simulation and X-Ray Scattering Study of Proteoliposomes

**DOI:** 10.64898/2026.07.03.736288

**Authors:** Enrico F. Semeraro, Ladislav Bartoš, Paulina Piller, Rahul Deb, Sandro Keller, Robert Vácha, Georg Pabst

## Abstract

Integral membrane proteins remodel the surrounding lipid bilayer, but quantifying the resulting deformations and linking them to protein density in the membrane has remained challenging. Here, we introduce an integrative methodology that combines all-atom molecular dynamics (MD) simulations with multiscale small-angle X-ray scattering (SAXS) analysis to connect membrane strain to the protein/lipid ratio in proteoliposomes. Using outer membrane phospholipase A (OmpLA) reconstituted into lipid bilayers with both increased and decreased hydrophobic thickness, we systematically probe the effects of positive and negative hydrophobic mismatch. MD simulations demonstrate that OmpLA causes anisotropic, oscillatory thickness deformations extending up to eight times the radius of the first lipid shell surrounding the protein, yet the net change in average membrane thickness remains below 1%. Through our multiscale SAXS analysis, we quantitatively extract structural parameters, ranging from proteoliposome size to internal membrane architecture, using constrained Bayesian inference, with priors derived from MD findings. Specifically, we determine the protein/lipid molar ratio and average membrane strain, revealing excellent agreement between experiment and simulation. In thinner bilayers, substantial protein loss limits the analysis, highlighting the role of bilayer stability in sample preparation. Moreover, the predominance of OmpLA monomers in the thicker membranes is consistent with weak, membrane-mediated repulsive interactions between protein inclusions. Collectively, this integrative approach establishes a framework for quantifying protein–lipid interactions across molecular and mesoscale dimensions.

## I. INTRODUCTION

Biological membranes are dynamic, compositionally complex assemblies of lipids and proteins that underpin key cellular functions and organizational principles.^1^ Far from being inert barriers, membranes serve as active platforms for signaling, substrate transport, and free-energy transduction, with their emergent properties arising from intricate physical and chemical interactions among their molecular constituents.^2^

A central challenge in membrane biophysics is to unravel how the interplay between lipids and integral membrane proteins governs both the architecture and the function of the bilayer. Protein–lipid interactions are not restricted to direct contacts, but propagate through the bilayer, mediating changes in protein conformation, organization, and activity via collective phenomena often described as membrane-mediated effects. For example, our recent work demonstrated that the enzymatic activity of a membrane protein is tuned by curvature stress differences across asymmetric bilayers, underscoring the profound impact of the physical membrane environment on protein function.^3,4^

In this study, we turn the focus to the reciprocal relationship: the influence of integral membrane proteins on the structural properties and elastic response of the lipid bilayer. A particularly relevant framework is the concept of hydrophobic matching, where a mismatch between the hydrophobic transmembrane span of a protein and the thickness of the surrounding lipid bilayer leads to free-energy penalties.^5^ The protein– lipid system accommodates mismatch by adjusting local bilayer thickness, lipid packing, or protein topology,^6,7^ with consequences for long-range protein–protein interactions.^8–10^ Continuum mechanical models have provided significant insights into these processes, relating deformation amplitudes and ranges to membrane elasticity and protein geometry.^8,11–13^ Coarse-grained molecular dynamics (MD) simulations have extended these findings, offering detailed predictions for how protein inclusions deform the bilayer and interact over long distances.^9,12,14^ However, a quantitative, experimentally validated link between protein density in the membrane and the resulting bilayer deformation has remained elusive. Existing evidence largely relies on NMR analyses of a small number of systems, often at non-physiologically low hydration levels.^10,15^

To address this gap, we present an integrative approach that combines high-resolution all-atom MD simulations with small-angle X-ray scattering (SAXS) experiments. SAXS enables robust, ensemble-averaged structural characterization of fully hydrated, free-floating unilamellar vesicles with defined lipid and protein composition. To accurately interpret the scattering data, we developed a multiscale SAXS analysis framework that connects global vesicle size and protein density to local membrane structure and organization. This approach leverages detailed models of lipid bilayer structure^17,18^ and Membrane Thickness Strain from Protein Inclusions adapts classical colloidal and polymer scattering methods for finite-sized, decorated vesicles.^19,20^ By bridging three orders of magnitude in length scale, our model provides a statistical ensemble describing both overall vesicle architecture and the local structural response of the membrane, including thickness, lipid packing, and protein-protein separation.

As a model system, we focused on the outer membrane phospholipase A (OmpLA), a well-characterized enzyme that hydrolyzes phospholipids upon dimerization but remains inactive in the absence of Ca^2+^, ensuring membrane integrity throughout the experiment.^21^ We reconstituted OmpLA into ∼100 nm unilamellar vesicles composed of either 1-palmitoyl-2-oleoyl-*sn*-glycero-3-phosphocholine (POPC) or 1,2-dilauroyl-*sn*-glycero-3-phosphocholine (DLPC), thus exploring the effects of positive hydrophobic mismatch (POPC, where the bilayer is thicker than the hydrophobic span of OmpLA) and negative hydrophobic mismatch (DLPC, where it is thinner).^7^

Our study reveals that OmpLA induces complex, anisotropic, long-range oscillatory deformations in the membrane, extending up to eight lipid shells from the protein boundary yet producing a net change in membrane thickness of less than 1%. Integrating MD simulations with SAXS, we quantitatively extract protein/lipid ratios and average membrane strain from scattering data, with results in excellent agreement between simulations and experiments. This approach establishes a generalizable framework for interrogating the multiscale coupling between proteins and lipid bilayers in both biological and synthetic membrane systems.

## II. MATERIALS AND METHODS

### A. Materials

POPC, DLPC, 1-palmitoyl-2-oleoyl-*sn*-glycero-3-phospho-(1’-*rac*-glycerol) (POPG), and 1,2-dilauroyl-*sn*-glycero-3-phospho-(1’-*rac*-glycerol) (DLPG) were purchased from Avanti Polar Lipids (Alabaster, AL, USA). Tris(hydroxy-methyl) aminomethane (Tris) was obtained from Carl Roth (Karlsruhe, Germany), lauryldimethylamine-*N*-oxide (LDAO) was acquired from Sigma-Aldrich (Vienna, Austria), and ethylenediaminetetraacetic acid (EDTA) from Merck (Darmstadt, Germany). The full list of chemicals and solvents for OmpLA production, purification and reconstitution are available in the Supplementary Material, Sec. S1.

### B. Preparation of proteoliposomes

Large unilamellar proteoliposomes with a size of about 100 nm (PLUVs) were prepared following established protocols.^3,22^ Briefly, purified OmpLA was refolded into LDAO micelles and reconstituted into extruded 100 nm-sized large unilamellar lipid vesicles (LUVs) by titration at targeted protein/lipid molar ratios, 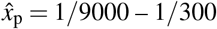. The LUVs were either composed of POPC, doped with 5 mol% POPG, or DLPC, doped with 5 mol% DLPG. At these low concentrations, the anionic lipid component does not significantly affect membrane structure or protein activity.^4^

LDAO was then removed by dialysis against 20 mM Tris, 2 mM EDTA buffer overnight, yielding PLUVs after extruding through a 100 nm filter. As controls, we also prepared *mock* lipid-only samples, which underwent the same preparation protocol as the protein-containing samples. Further details regarding the preparation of these samples can be found in the Supplementary Material, Sec. S1.

The concentration of reconstituted OmpLA was determined using UV-Vis spectroscopy with a NanoDrop ND-1000 spectrophotometer (Peqlab Biotechnology GmbH, Erlangen, Germany), while the final lipid concentration was quantified through a phosphate assay. Both methods are described in detail in a previous study.^3^ Notably, the apparent protein/lipid molar ratio, *x*_p_, was consistently larger than the targeted 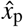, for both POPC and DLPC systems (Fig. 1). This observation indicates a net loss of lipid relative to protein during the reconstitution process.

**FIG. 1.**
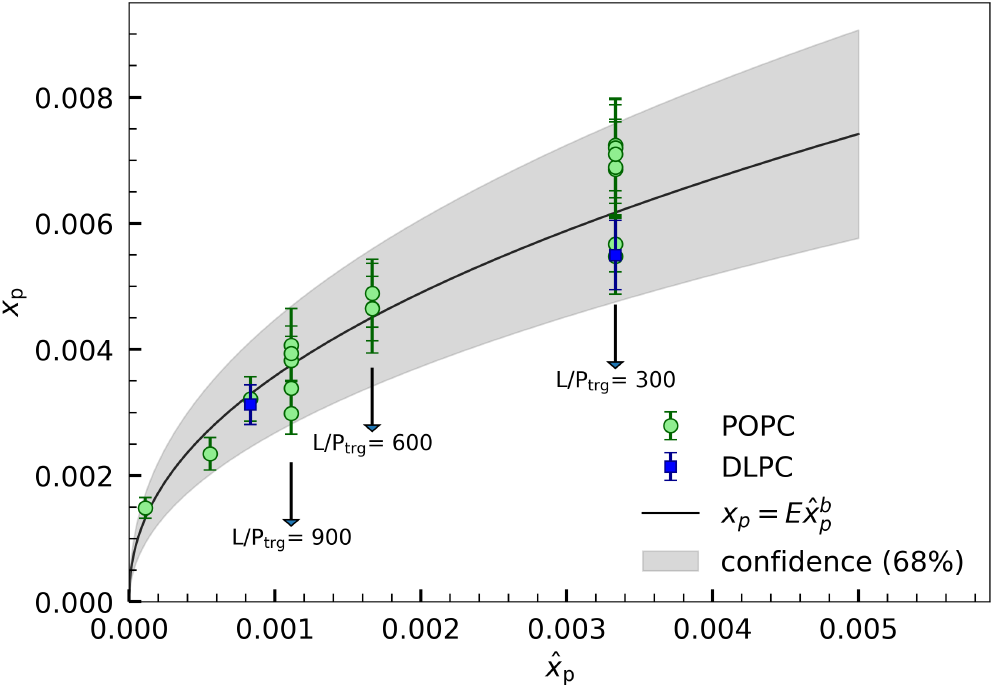
Comparison between targeted and apparent protein/lipid molar ratios 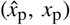 for OmpLA in POPC (green circles) and DLPC (blue squares) vesicles. Data were fitted with *x*_p_ = (0.082 ± 0.014) · 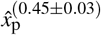 (black line). The gray area represents the 68% confidence range.

### C. SAXS experiments

POPC based samples (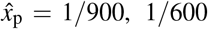, and and 1/300) were measured at the BioSAXS beamline P12 operated by EMBL Hamburg at the PETRA III storage ring (DESY, Hamburg, Germany). Samples were measured at a concentration of ∼ 7 mg/ml at 37 ^°^C in a flow-through quartz capillary with a diameter of 1 mm. Data were acquired with the beamline standard setup, and automatic data reduction was performed on site https://www.embl.org/groups/small-angle-x-ray-scattering/.^23^

DLPC based samples (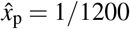, and 1/300) were measured with a SAXSpoint 5.0 camera (Anton Paar, Graz, Austria) equipped with a MetalJet X-ray generator (Excillum, Kista, Sweden) and an Eiger R 1 M detector system (Dectris, Baden-Dättwil, Switzerland). Data was integrated and background subtracted using SAXSanalyis (Anton Paar). Samples were measured at a concentration of ∼ 20 mg/ml at 37 ^°^C using the *µ*Cell (Anton Paar).

### D. Molecular dynamics simulations

All-atom simulations were performed with GROMACS 2021.4^24^ using the CHARMM36m force field^25,26^ for lipids and proteins and TIP3P water.^27^ The monomer and dimer structures were obtained from the Protein Data Bank (PDB IDs 1QD5 and 1QD6, respectively).^21^ The missing residues of the dimer structure were modeled using the coordinates of the monomer structure. The protein structures were minimized in vacuum using the steepest descent algorithm.

#### a. Systems preparation

Flat membranes composed of POPC or DLPC bilayers with embedded monomeric or dimeric OmpLA were prepared using the CHARMMGUI.^28,29^ To investigate the effects of the protein/lipid molar ratio (*x*_p_), we simulated two regimes: low-*x*_p_ and high-*x*_p_.

In the low-*x*_p_ regime, a single protein unit was embedded in a large membrane patch to ensure the bilayer could relax to its protein-free, bulk thickness. This configuration minimizes protein crowding and enables the membrane to maintain properties close to those of pure lipid bilayers. Specifically, simulations were performed with 800 lipids for systems containing monomeric OmpLA and 1200 lipids for dimeric OmpLA.

In the high-*x*_p_ regime, system size and composition were chosen to mimic the highest experimental protein/lipid ratios measured in PLUVs (see Fig. 1). Here, the average protein-protein distance along the vesicle surface was estimated as

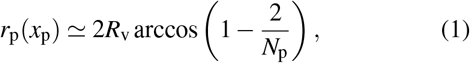

where *N*_p_ is the number of proteins per vesicle and is defined by

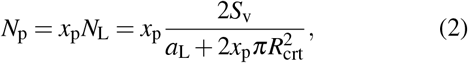

with *N*_L_ as the average number of lipids per PLUV and 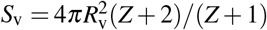 representing the average vesicle surface area. Here, *Z* is the width parameter of the Schultz distribution function used to describe the polydispersity in the vesicle radius, *R*_v_ (see B), *a*_L_ = *V*_HC_*/D*_C_ is the average area per lipid, and *R*_crt_ is the cylindrical radius used to approximate the inplane protein shape (see Section III). *V*_HC_ and *D*_C_ denote the acyl-chain volume and half the hydrophobic membrane thickness, respectively.

Based on these relationships and using parameters of *a*_L_∼ 66^2^, *R*_v_ = 450, and *Z* = 10, we constructed simulation boxes corresponding to high-*x*_p_ regimes of approximately 85 for monomeric and 130 for dimeric OmpLA. This translated to systems containing approximately 200 lipids for monomers and 500 lipids for dimers.

#### b. Simulation

All systems were solvated with water and contained NaCl ions at physiological concentrations, with additional ions added to ensure overall charge neutrality. The constructed systems were energy-minimized using the steepest-descent algorithm with positional restraints on the protein atoms. Subsequently, each system was equilibrated in six sequential steps, with progressively increasing simulation lengths and time steps. The first two stages were performed in the canonical (NVT) ensemble, while the subsequent stages were conducted in the isothermal-isobaric (NPT) ensemble. The temperature was kept at 310 K, with three separate thermal baths assigned to the protein, membrane, and water with ions. During the NPT equilibration stages, the pressure was maintained at 1 bar with a semi-isotropic coupling scheme. The positional and dihedral restraints applied during the minimization step were also used during equilibration, with their strengths gradually reduced over the course of the equilibration process. Following equilibration, each system was simulated in three independent replicas for a total of 2 *µ*s per replica. The simulation parameters were identical to those used in the final stage of equilibration, with two key exceptions: the Berendsen barostat was replaced by the Parrinello–Rahman barostat,^30,31^ and all positional and dihedral restraints were removed. Further details regarding the minimization, equilibration, and production protocol are provided in the Supplementary Material, Sec. S2.

#### c. Analysis

The final 1.5 *µ*s of each simulation trajectory were analyzed to determine membrane properties. Membrane thickness, defined as the phosphate-to-phosphate distance (*D*_pp_), was calculated using our in-house tool memthick (github.com/Ladme/memdian). For each replica, two-dimensional thickness maps were generated by dividing each membrane leaflet into 0.1 × 0.1 nm^2^ grid cells and computing the average *z*-position of the phosphate beads. The thickness in each bin was then calculated as the difference between the average *z*-positions of the phosphates in the upper and lower leaflets.

These 2D maps were further used to assess membrane strain by comparing the local acyl-chain thickness *D*_C_(**r**) to its bulk value 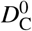 :

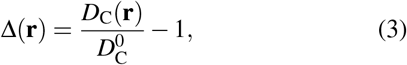

where **r** is the in-plane distance from the protein’s center of mass (CoM). *D*_C_ was estimated from *D*_pp_*/*2 by subtracting 4.2 Å, which accounts for the average distance from the lipid backbone to the phosphate group.^18,32^

One-dimensional profiles of membrane thickness were then obtained from the 2D maps using two methods. In the low-*x*_p_ regime, radial averaging was performed around the central OmpLA protein, producing a thickness profile as a function of distance from the protein. In the high-*x*_p_ regime, the periodic simulation box was expanded to include proteins from neighboring replicas, and thickness was averaged along lines connecting the central protein unit to its four nearest virtual neighbors. In both cases, the resulting 1D profiles were converted to deformation profiles as described above. Finally, Membrane Thickness Strain from Protein Inclusions profiles from individual replicas were averaged, with the standard deviation providing an estimate of the associated error.

## III. MULTISCALE SCATTERING MODEL

We propose a multiscale model for the analysis of SAXS data from PLUVs. The model provides key structural information, including vesicle size, transbilayer structure, and the conformation, density (i.e., in-membrane concentration), and spatial distribution of integral membrane proteins. A notable advantage of this model is its modularity, which allows contributions from the vesicle and the protein to be modeled independently.

For a single PLUV the scattering intensity can be expressed as:

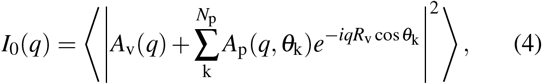

where *q* is the modulus of the scattering vector, *A*_v_(*q*) is the scattering amplitude of a single vesicle, *A*_p_(*q*) is the scattering amplitude of a single integral membrane protein, *N*_p_ is the average copy number of proteins per vesicle, and *θ*_k_ is the polar angle defining the position of the *k*-th protein within the spherical frame of the vesicle of radius *R*_v_ (Fig. 3A). *R*_v_ coincides with the center of the lipid bilayer as well as the radial center of mass (CoM) of the proteins. Small displacements of the proteins from this position (on the order of a few angstroms) are negligible compared to the vesicle size. The brackets ⟨· · · ⟩ denote ensemble averaging over the entire population of PLUVs.

For a suspension of non-interacting PLUVs with number density *n*_v_, solving Equation 4 yields

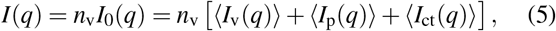

where ⟨*I*_v_(*q*) ⟩, ⟨*I*_p_(*q*) ⟩, and ⟨*I*_ct_(*q*⟩ denote the average scattering intensities arising from the vesicles, the proteins, and the vesicle-protein cross-term, respectively. Figure 2 displays the individual contributions of each component to the total scattering intensity. At low *q, I*(*q*) is dominated by ⟨*I*_p_(*q*) ⟩, while ⟨*I*_v_(*q*) ⟩ becomes dominant at high *q*. The cross-term ⟨*I*_ct_(*q*)⟩ changes sign near the first minimum of ⟨*I*_v_(*q*) ⟩ and influences *I*(*q*) across all *q* values.

**FIG. 2.**
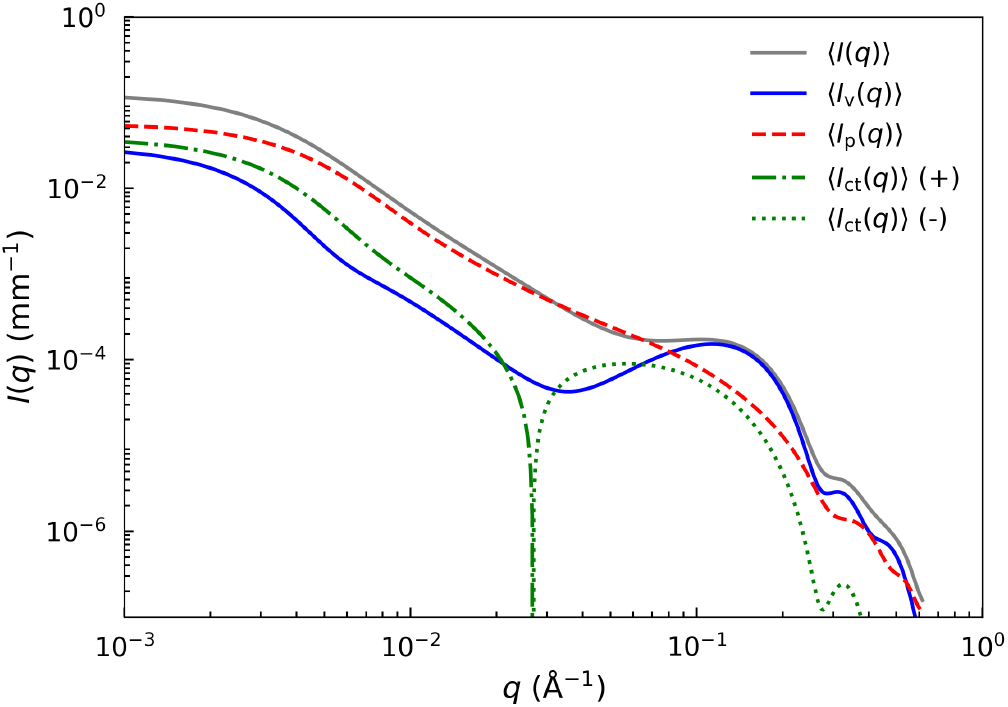
Calculated scattering intensity of OmpLA monomers in POPC vesicles (*x*_p_∼ 0.006) using Equation 5, along with its individual components: vesicle contribution (⟨*I*_v_⟩, solid blue line), protein contribution (⟨*I*_p_⟩, dashed red line), and the vesicle-protein cross-term (⟨*I*_ct_ ⟩, dashed-dotted green line for positive values and dotted green line for negative values).

### A. Vesicle term

For analyzing lipid vesicle scattering, the most detailed approach combines two models: the scattering density profile (SDP) model for lipid bilayers^17,18^ and the separated formfactor (SFF) model.^33^ This hybrid SFF-SDP model enables a comprehensive characterization by treating the bilayer structure and vesicle size effects separately. The contribution from vesicles to *I*(*q*) then becomes

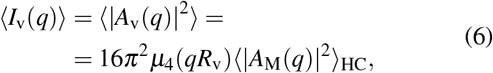

where *A*_M_(*q*) is the one-dimensional scattering amplitude of a planar lipid bilayer with a normal distribution of the hydrocarbon thickness ⟨· · · ⟩ _HC_^18^ (see Appendix A for details).

The term 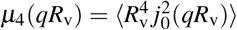 accounts for polydispersity in vesicle sizes. Here, *j*_0_(*x*) is the 0^th^-order spherical Bessel function of the first kind (see Appendix B).

### B. Protein term

The protein term captures both the structural details of individual membrane proteins (*form factor*), and their number density and spatial arrangement on the vesicle surface (*structure factor*).

#### 1. Protein form factor

We first derive an analytical representation of the scattering amplitudes of bare OmpLA embedded in POPC or DLPC membranes. To this end, we take the final snapshot from each MD-equilibrated OmpLA structure and remove all contributions from the membrane and aqueous solvent. We then compute the corresponding vacuum scattering form factors using CRYSOL^34^ and verify that the resulting scattering intensities exhibit no significant configuration-dependent discrepancies (Fig. S3A–B). From these data, we obtain ensemble-averaged vacuum scattering intensities for OmpLA monomers and dimers (Fig. S3C). Finally, we replace the physical protein with a *virtual protein* model that reproduces the same scattering intensity while providing a simplified analytical form for the scattering amplitude (Fig. 3B).

The chosen virtual protein model is a stack of homogeneous coaxial disks with the scattering amplitude

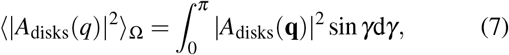

where ⟨… ⟩ _Ω_ is the orientational average, *q*_∥_= *q* cos(*γ*) and *q*^⊥^ = *q* sin(*γ*). For details, see Appendix C. The disk parameters (size, thickness, number) are adjusted until good agreement of the scattering intensity with the CRYSOL data is achieved (Fig. 3C, Table V).

For vesicle insertion (Fig. 3D), we first add a hydration layer to both luminal and extravesicular protein surfaces. The scattering amplitude of the hydrated virtual protein is

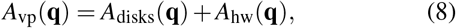

where *A*_hw_(**q**) refers to the hydration layer (thickness *d*_hw_ = 3.1 Å, SLD *ρ*_hw_ = 11.5 × 10^−6^ Å^−2^, see^35^ and C7).

**FIG. 3.**
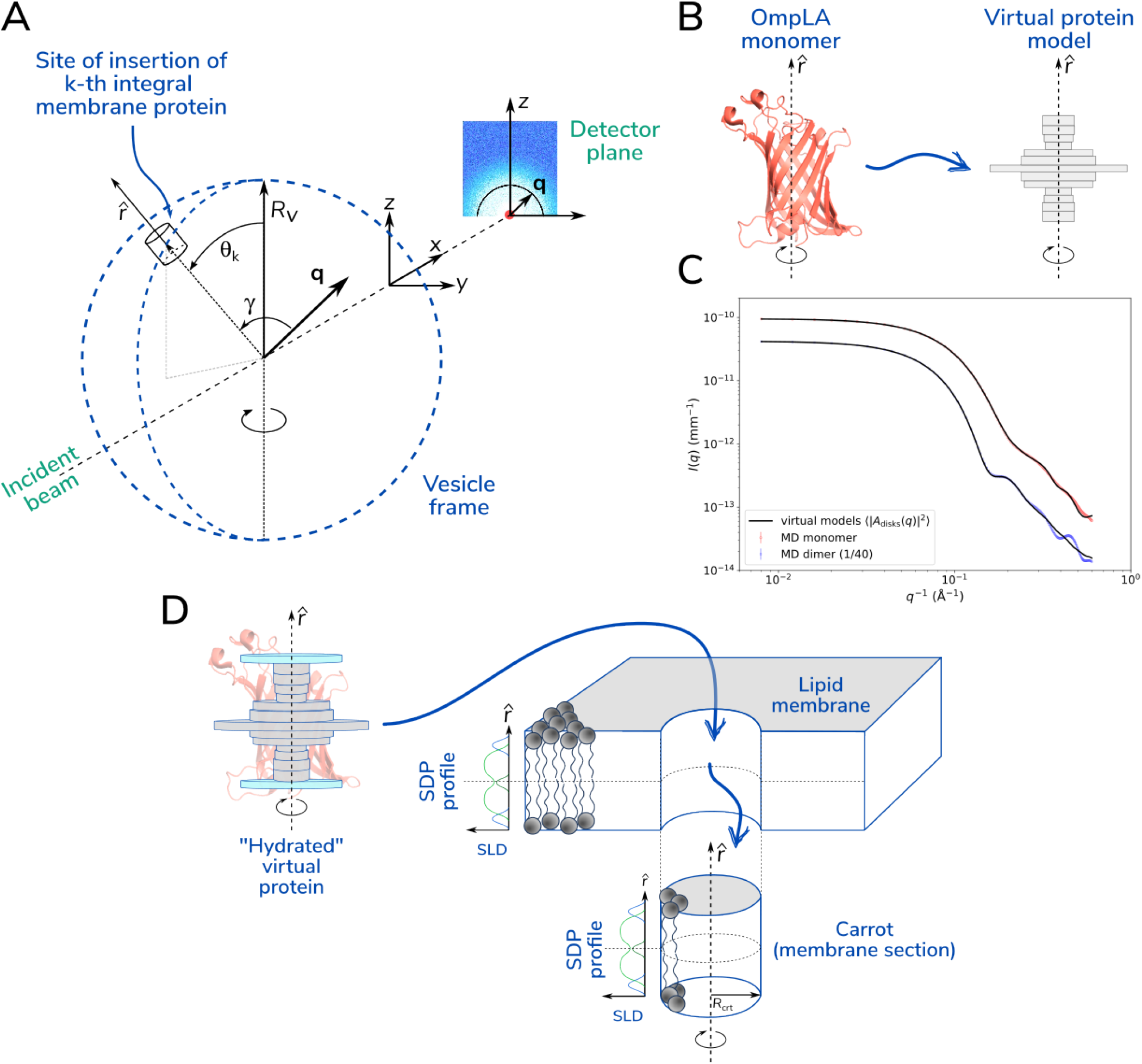
Overview of geometries and definitions used in the multiscale model. A) Vesicle frame of radius *R*_v_ with protein inclusions. The location of the k^th^ protein on the spherical vesicle is given by the polar angle *θ*_k_, where the protein director 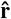 aligns normal to the vesicle surface, and 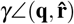 Insertion of the virtual protein into the lipid bilayer requires removal of an equivalent cylindrical membrane section of radius *R*_crt_.

Next, to model protein insertion, we subtract the bilayer volume occupied by the protein, approximated as a cylinder (the “carrot”, *R*_crt_ = 17.2 Å for monomers, 20.3 Å for dimers). The carrot’s scattering amplitude is

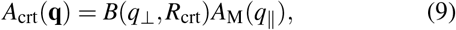

where *B*(*q*^⊥^, *R*_crt_) is the cylinder’s radial amplitude and *A*_M_(*q*^∥^ ) is the SDP-based membrane amplitude parallel to the protein director (see Appendix C and Fig. 3A, C).

The final form factor for a membrane-embedded protein becomes

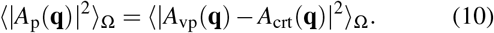

#### 2. Protein structure factor

To model the spatial distribution of the proteins within vesicles, we adapted the analytical framework developed for spherical micelles of block copolymers, Pickering emulsions and raspberry particles.^19,20^ A key assumption of this method is the homogeneous distribution of ‘decorating entities’ on the particle surface, that leads to the effective protein structure factor

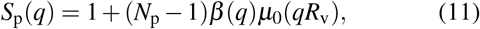

where *µ*_0_(*qR*_v_) is the 0^th^ raw moment of 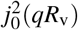accounting for the *R*_v_-polydispersity (Appendix B). The term *β* (*q*), is a second-order correction that accounts for the orientation distribution of anisometric objects^36^ (see Appendix D for details), and is expressed as

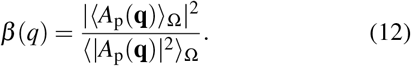

Finally, combining Eqs. (10) and (11), the full protein term in Eq. (5) is given by

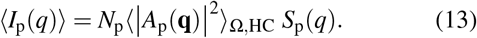

### C. Vesicle-protein cross term

Following the method in^19^, the vesicle–protein cross-term in the scattering intensity is given by

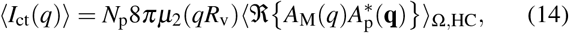

where *µ*_2_(*qR*_v_) is the second raw moment of 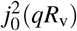 (see Appendix B). The notation ℜ(· · ·) indicates that only the real part of the expression inside the brackets is taken. Importantly, *A*_M_(*q*) and *A*_p_(**q**) are complex functions only if the bilayer or the virtual protein models are asymmetric.^18,37^

### D. Protein/lipid ratio

The forward scattering at *q* → 0 for the single-PLUV form factor (Eq. 4) is given by

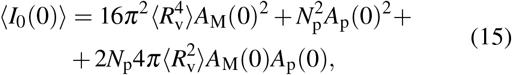

where

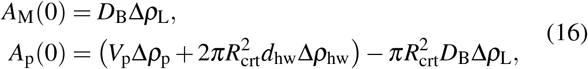

with *D*_B_ representing the Luzzati thickness (the total thickness of the lipid bilayer), and Δ*ρ*_L_ the contrast between the average scattering length density (SLD) of the membrane and that of the solvent.

This expression for the forward scattering ensures that the lipid and protein contributions are properly balanced, making it possible to quantitatively extract the value of 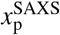 from experimental scattering data via Eq. (2). When 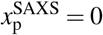, the model naturally reduces to the standard LUV model.^18^

## IV. RESULTS

### A. Short- and long-range membrane thickness strain

To elucidate the membrane perturbations induced by OmpLA, we conducted MD simulations of both OmpLA monomers and dimers embedded in flat bilayers, examining both the low-*x*_p_ and high-*x*_p_ regimes as detailed in Section II D.

Analysis of in-plane thickness changes, represented by Δ(**r**) (Eq. 3), reveals a relatively uniform zone of membrane thinning (Δ(**r**) *<* 0) surrounding OmpLA monomers in POPC bilayers; this effect is consistently observed in both low- and high-*x*_p_ conditions (Fig. 4A, C). This thinning region is followed by a ring of increased bilayer thickness (Δ(**r**) *>* 0), with the membrane relaxing back to its bulk thickness only in the low-*x*_p_ regime.

**FIG. 4.**
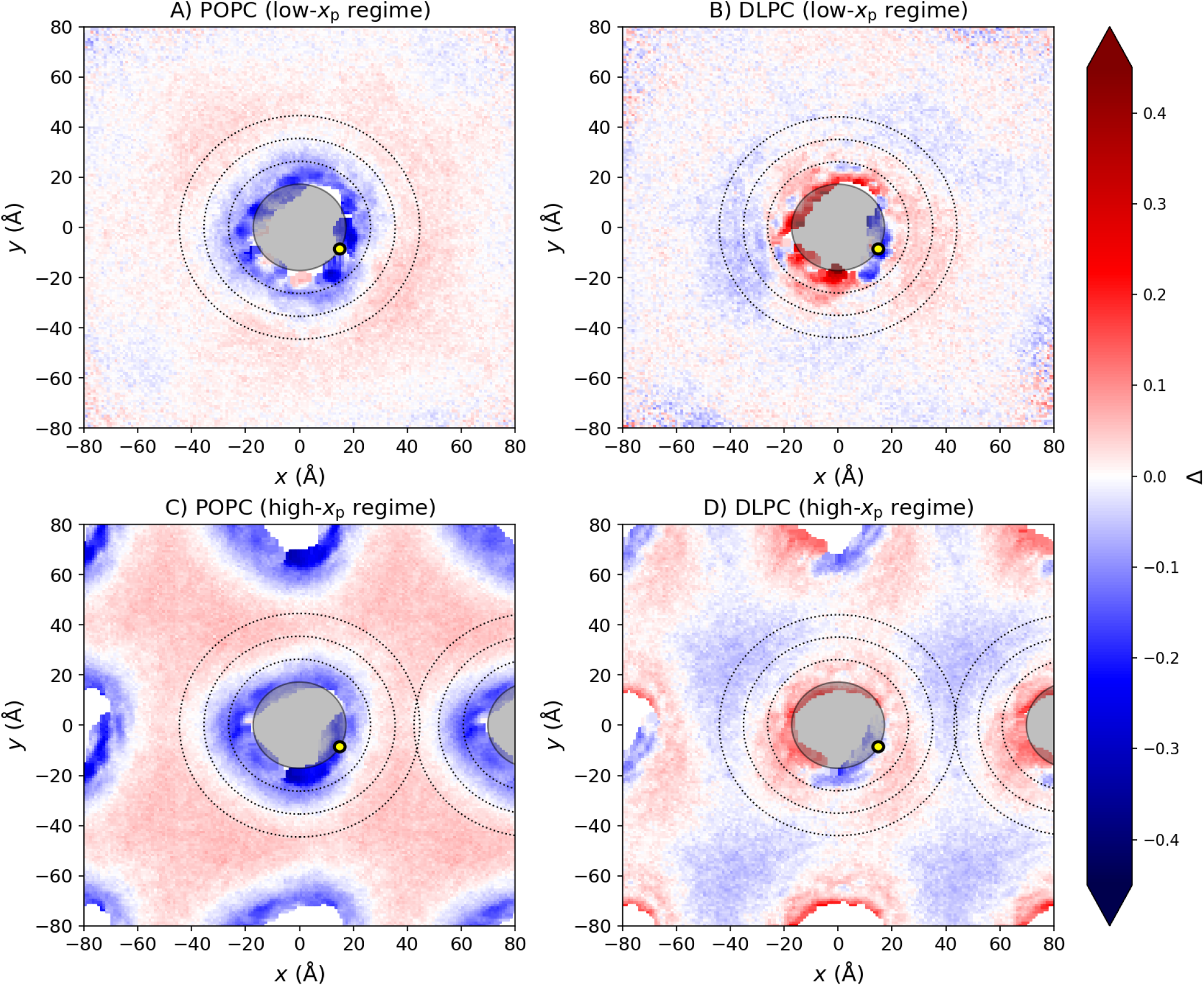
Relative lipid thickness strain, Δ, around OmpLA monomers. Panels **A-B** depict isolated OmpLA in the low-*x*_p_ regime, while panels **C-D** represent the crowded high-*x*_p_ regime. The color scale illustrates relative membrane thickening (red) and thinning (blue). The shaded circle corresponds to the protein’s cross-section with radius *R*_crt_, and the dotted lines indicate the average boundaries of the first three lipid layers. For reference, the yellow dot marks the location of the protein’s active site.

The membrane response in DLPC bilayers differs markedly, as illustrated in Fig. 4B and D. Here, the perturbation is more heterogeneous, showing areas of both localized thinning and thickening in the vicinity of OmpLA. Comparable membrane deformation patterns are observed for OmpLA dimers (see Fig. S1), where the bilayer responds similarly to the monomer systems.

The resulting strain profiles display oscillatory behavior around both OmpLA monomers (Fig. 5) and dimers (Fig. S2), with membrane strain reaching up to 15% near the protein surface. Importantly, local membrane strain extends well beyond the first lipid shell (defined as 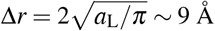, for *a*_L_ ∼ 66 Å^2^), and is more pronounced in POPC than in DLPC, and for dimers compared to monomers. In the high-*x*_p_ regime, the bilayer does not fully recover its bulk thickness, indicating persistent membrane perturbation under crowded conditions.

**FIG. 5.**
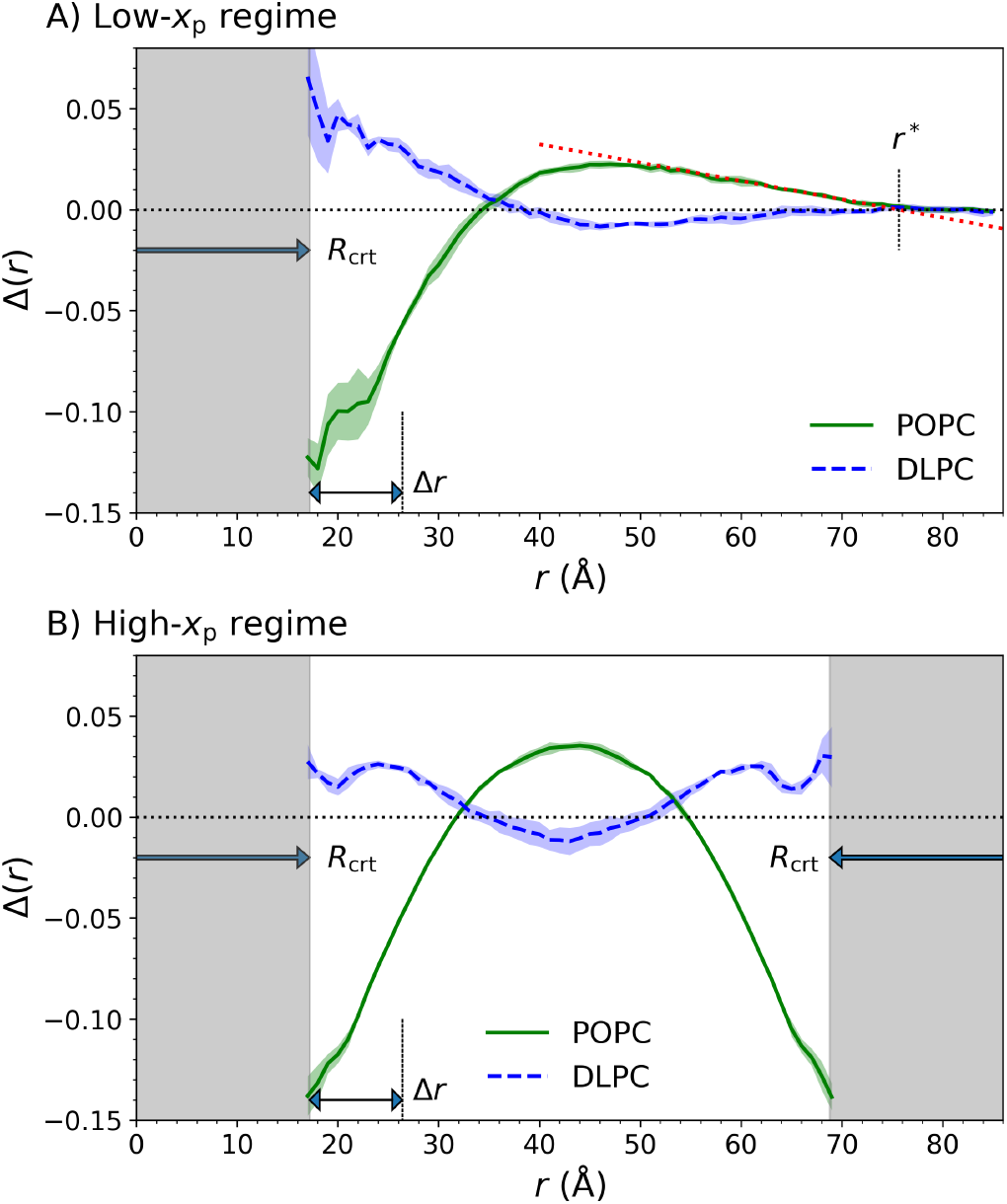
Average membrane strain profiles in POPC (solid green) and DLPC (dashed blue) bilayers for OmpLA monomers, with the protein region indicated by gray shading. Each curve represents the average over three independent replicas, and the shaded bands denote the corresponding standard error envelopes. Δ*r* marks the extent of the first lipid shell surrounding the protein. **(A)** Low-*x*_p_ regime: The dotted red line illustrates the determination of *r*^∗^ for POPC. **(B)** High-*x*_p_ regime: Averaging is performed along a line connecting two OmpLA inclusions.

To quantify the spatial extent of membrane perturbations in low-*x*_p_ systems (Fig. 5A), we defined the radius of the deformation zone, *r*^∗^, as the intersection between the baseline and the tangent to Δ(*r*) for *r < r*^∗^. The average number of lipids affected within this zone is given by:

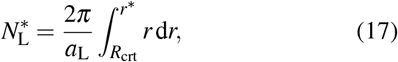

while the mean membrane thickness strain *per OmpLA* monomer (or dimer) is expressed as

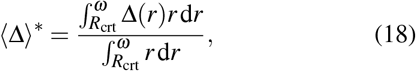

where *ω* is set to *r*^∗^ in the low-*x*_p_ regime, and *r*_p_*/*2 for high-*x*_p_. Our analysis reveals that the perturbation zone can reach up to ten times the radius of the first lipid shell, encompassing several hundred lipids. Despite this significant spatial range, the average membrane thickness strain remains small (Tab. I). In most cases, ⟨Δ⟩^∗^ values are indistinguishable from zero, reflecting the oscillatory character of the deformation profiles, in which regions of local thinning and thickening largely cancel upon area-weighted averaging.

### B. Counting proteins and the effective membrane strain in PLUVs

SAXS data analysis was performed using the multiscale scattering model (Eq. 4), accounting for OmpLA as either monomers or dimers. Fits for monomers appear in Fig. 6, with dimer fits in Fig. S4.

**FIG. 6.**
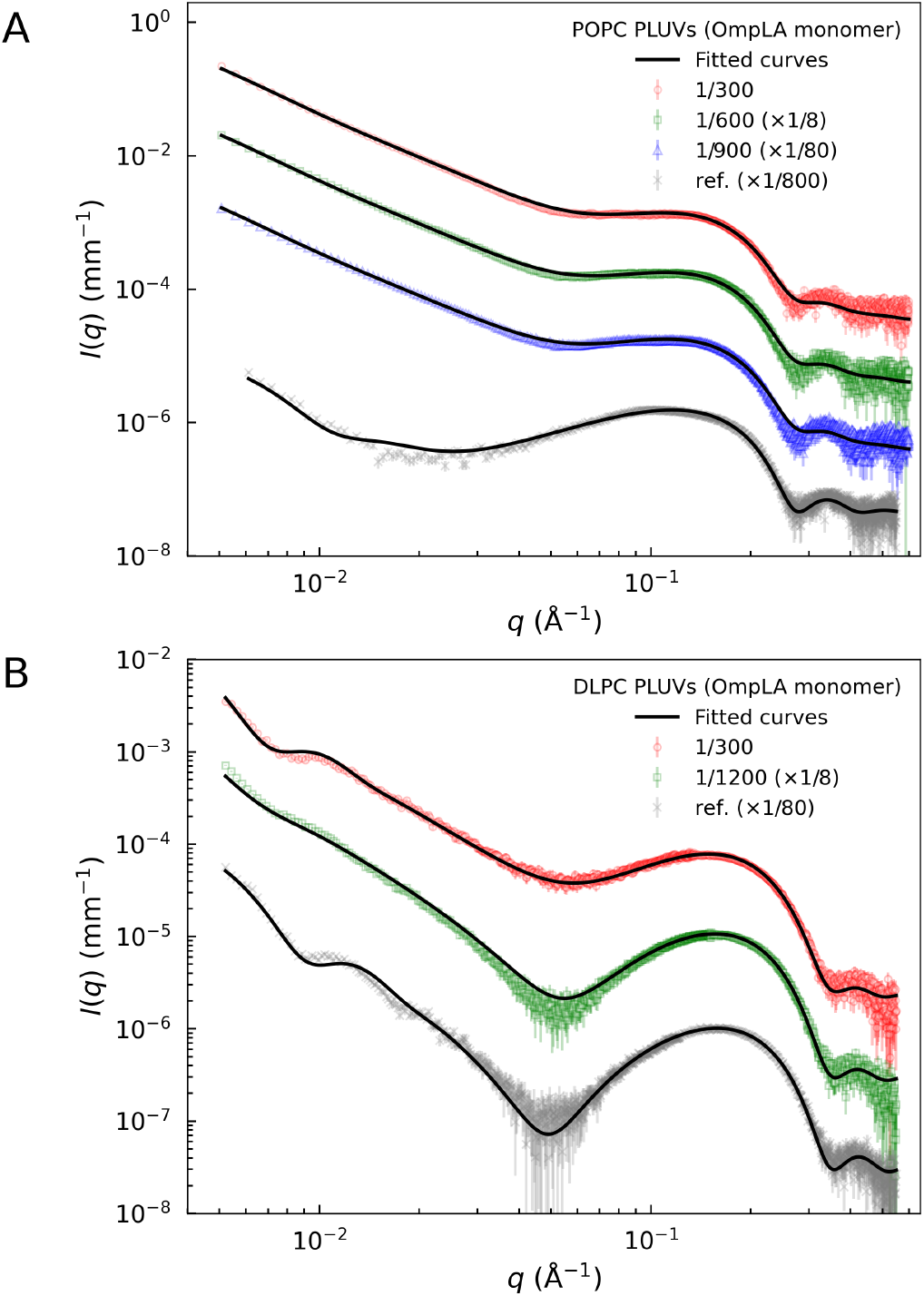
SAXS data and fitted curves of POPC (A) and DLPC (B) PLUVs at different 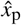 conditions in the case of OmpLA monomers (data and fits were scaled to improve visibility). References curves refer to *mock* lipid-only samples.

All analyses used the *SAS_MoCa* toolkit,^38,39^ a stochastic minimization algorithm that exploits constrained Bayesian inference to address the overparameterization of the SDP model. This method incorporates informative Gaussian priors to guide fitting and improve convergence towards the reduced chi-squared, *χ*^2^, global minimum. Further details on prior selection can be found in the Supplementary Material, Sec. S4 and Tabs. S2–S4.

Analysis of goodness-of-fit, assessed via *χ*^2^ values (Fig. S5), strongly indicates that OmpLA is predominantly monomeric in POPC membranes. This is further supported by protein concentration-dependent changes in the water volume of the lipid hydration layer. Models assuming OmpLA dimers result in unphysical fits under these conditions

(Fig. S8). Specifically, the fitted volume of water molecules in the lipid hydration shell, *V*_BW_, decreases systematically with increasing 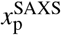 in the monomer model, consistent with the partial overlap of annular lipids with the hydration shell of the adjacent protein. In contrast, the dimer model drives *V*_BW_ against the boundaries of the prior distribution, indicating a non physical underestimation of water volume in the hydration shell (Sec. S4).

For DLPC membranes, the apparent amount of OmpLA incorporated was very low, making it difficult to distinguish between monomeric and dimeric states. Complete parameter fitting results are compiled in Supplementary Tables S5–S8.

Figure 7 compares protein/lipid molar ratios determined by SAXS 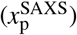 with those obtained from optical methods (Fig. 1). In POPC, the two techniques agree closely 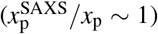. However, for DLPC, 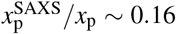, indicating about sixfold lower protein concentration in PLUVs. Using the fitted values for 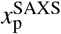, *R*_v_, *Z* and *a*_L_ (Tables S5– S8), we calculated *N*_p_ with Eq. (2). The critical protein concentration 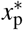, which marks the transition between low- and high-*x*_p_ regimes, was obtained by substituting 2*r*^∗^ into the inverse of Eq. 1. This analysis yields *N*_p_ values of 230–300 for POPC and less than 100 for DLPC. With 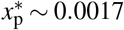 for both lipid types, these results confirm that POPC-based PLUVs are in the high-*x*_p_ regime, while DLPC systems are placed in the low-*x*_p_ regime (Fig. 7).

**FIG. 7.**
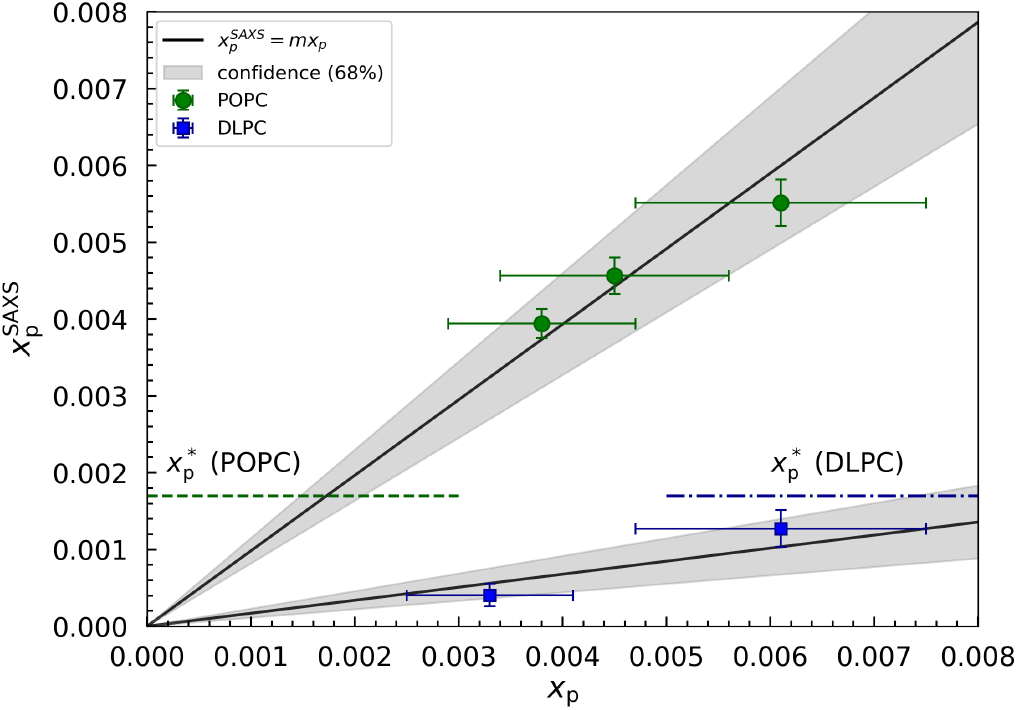
SAXS analysis results for 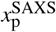 as a function of the effective *x*_p_ values, assuming a population of OmpLA monomers. Black line indicate the best fit or 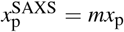, while the gray area represents the margin of error at the 68% confidence level. POPC: *m* = 0.98 ±0.17, DLPC: *m* = 0.17 ±0.06. Dashed and and dotted-dashed horizontal lines mark the transition from low to high-*x*_p_ regimes for POPC and DLPC, respectively.

Among the parameters obtained from our multiscale SAXS analysis, the hydrocarbon layer thickness can be tracked as a function of protein concentration. Using these data, we calculated the average membrane thickness strain, ⟨Δ ⟩^SAXS^, by constraining the analysis with the ⟨Δ⟩^∗^-values derived from MD simulations (Tab. I). To this end we apply

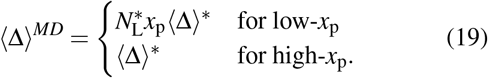

Figure 8 compares the experimental results with values from MD simulations across different 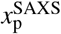, focusing on OmpLA monomers (with results for dimers shown in Fig. S7). Within experimental uncertainty, the agreement between SAXS and simulation is excellent, confirming that the average membrane thickness strain induced by OmpLA is small.

**FIG. 8.**
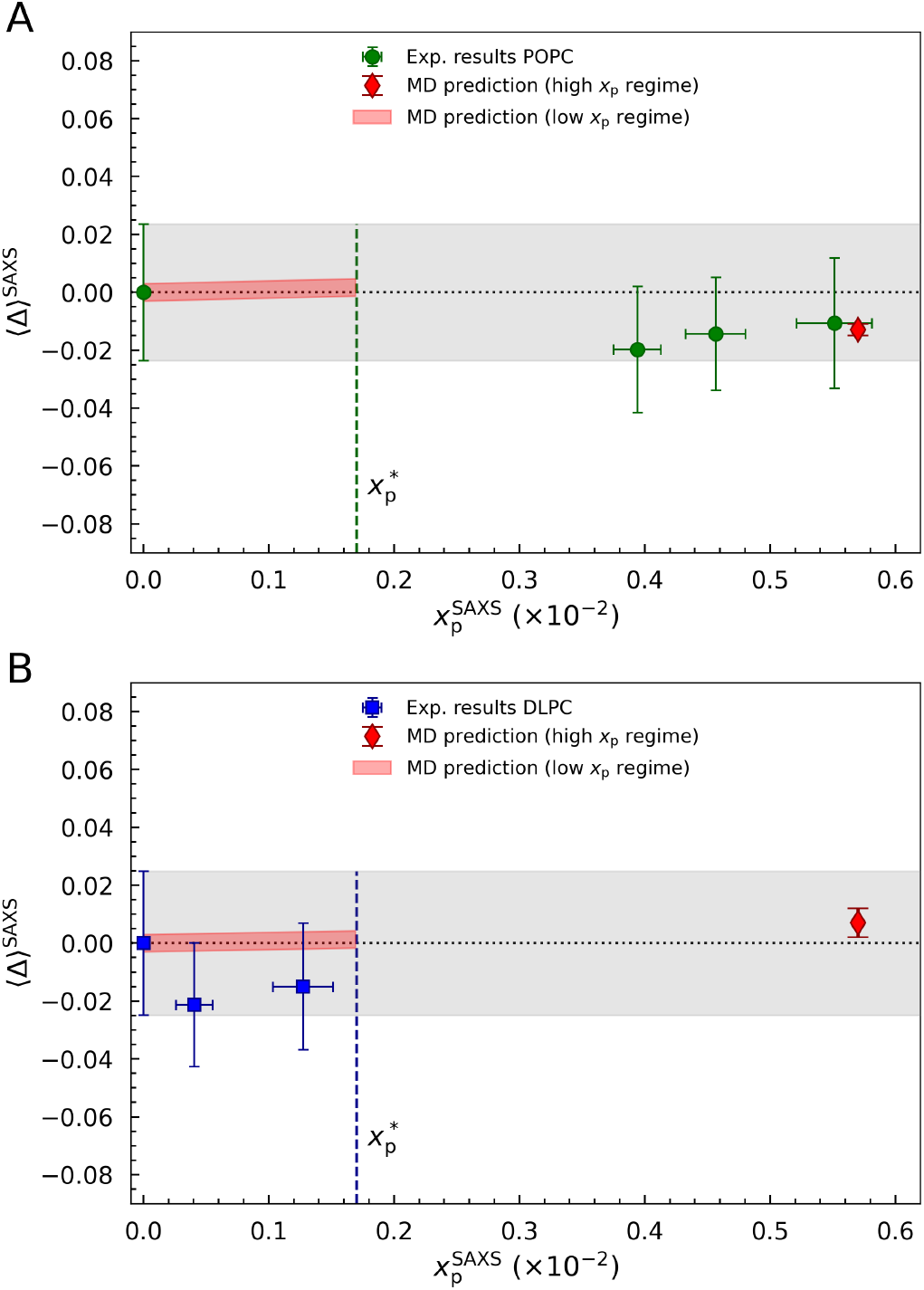
Average membrane strain, ⟨Δ⟩^SAXS^, for POPC (A) and DLPC (B) systems in the case of OmpLA monomers as a function of 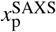. Dotted lines and gray areas represent reference values for bulk lipids and associated standard deviation. The predicted strain, ⟨Δ ⟩^MD^ is shown as red diamonds and red areas for the high-*x*_p_ and low-*x*_p_ regime, respectively. Vertical dashed lines highlight the transition from low to high-*x*_p_ regime.

## V. DISCUSSION

We show that integrating MD simulations with multiscale modeling of SAXS data yields a unified and quantitative structural description of ∼ 100 nm proteoliposomes (PLUVs). By directly connecting transbilayer lipid organization to the apparent protein/lipid molar ratio, our approach bridges the gap between molecular-scale interactions and experimentally accessible mesoscopic parameters. Importantly, our SAXS model does not aim to resolve fine structural details of the proteins, such as conformational changes or tilting, which are better addressed by complementary techniques.^40–42^

All-atom MD simulations reveal how OmpLA monomers and dimers deform membranes, capturing the effects of both low and high protein loadings. The multiscale SAXS model complements these findings by retrieving key structural features, including PLUV size distributions, average membrane thickness, and the apparent protein/lipid ratio. Importantly, the simultaneous determination of vesicle size and protein concentration allows direct comparison of MD predictions with SAXS-derived values, providing a robust framework for validating and interpreting results.

We distinguished three protein/lipid ratios: the targeted ratio 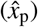, the optically measured apparent ratio (*x*_p_), and the SAXS-derived ratio 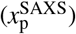. Consistently, we observe that 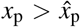, indicating a systematic loss of lipids during sample preparation. While the precise mechanism underlying this lipid depletion is not yet clear, the empirical power law dependence of *x*_p_ on 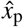 offers practical guidelines for future PLUV formulations.

Interestingly, in POPC PLUVs, *x*_p_ and 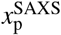 are in close agreement, with 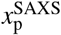 exhibiting 2–4 fold lower relative uncertainty. Conversely, for DLPC PLUVs, *x*_p_ exceeds 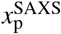 suggesting that protein losses may occur after PLUV formation but prior to SAXS measurement. Given that DLPC is the shortest disaturated phosphatidylcholine capable of forming bilayers^43^ it is less stable than POPC—particularly at elevated protein loading. This likely contributes to the formation of protein––lipid aggregates that are not detected by the used SAXS setup. This interpretation is consistent with earlier observations that OmpLA refolding into DLPC bilayers is inherently less efficient than into thicker membranes, with yields of only ∼ 40% in pure DLPC LUVs,^22^ likely owing to the higher prevalence of transient bilayer defects in these thin membranes.

We explored two MD simulation scenarios to probe the membrane response to OmpLA proteins. In the low-*x*_p_ regime, we examined membranes containing a single OmpLA monomer or dimer to assess how an otherwise bulk-like bilayer accommodates individual protein inclusions. For both POPC and DLPC bilayers, and for both protein oligomerization states, we observed damped, oscillatory thickness profiles surrounding the protein. Notably, these membrane deformations were more extensive around OmpLA dimers, with the affected zone reaching up to ten times the radius of the immediate lipid shell surrounding the proteins, even though the average protein-induced membrane strain remained relatively small. Interestingly, the SAXS analysis reveals a substantial increase in the hydrocarbon thickness fluctuation parameter, *ε*_C_, from ∼0.08 in protein-free membranes to ∼0.14 in POPC PLUVs (Tables S5). This broadening likely reflects the spatial heterogeneity in membrane thickness predicted by the MD simulations, where oscillatory deformation profiles create a distribution of local thicknesses that is averaged over the SAXS ensemble.

This oscillatory membrane response qualitatively matches previous results from coarse-grained simulations using idealized cylindrical protein models,^9^ and is consistent with predictions from continuum mechanical theories.^8,13^ The continuum models relate the amplitude and range of such deformations to bilayer elastic properties—such as bending and stretching moduli, spontaneous curvature, and interfacial mismatch. They predict that even modest, spatially extended deformations can significantly influence lipid-mediated protein–protein interactions, often in ways that remain experimentally elusive^8^. Specifically, the resulting membrane-mediated interactions can acquire a repulsive component, which may stabilize dispersed, non-aggregated protein distributions within the membrane.^11^

Beyond offering all-atom resolution, our study is distinguished by the use of a real, structurally complex protein with a 4–10-fold lower hydrophobic mismatch compared to previous models.^8^ The inherent inhomogeneity of the protein shape leads to heterogeneous and anisotropic deformation zones, which in turn shape local lipid–protein interactions at the protein surface. For example, in DLPC bilayers, while most annular lipids experienced positive membrane strain, a significant fraction instead experienced local bilayer thinning (see Fig. 4B). This spatial heterogeneity may increase the propensity for lipid scrambling at specific protein–lipid interfaces,^44^ supporting the emerging view that integral membrane proteins can dynamically modulate their immediate lipid environment.^2^

In the high-*x*_p_ regime, the membrane deformations induced by neighboring protein inclusions overlap significantly, effectively eliminating any regions of bulk-like lipid. This scenario supports previous MD findings, which showed that approximately 50–100 lipids co-diffuse with voltage-gated ion channels^45^, and, together with the observation that proteins occupy about 30% of the surface area in native membranes^46^, suggests that truly bulk lipids are largely absent in biological membranes. Our results not only reinforce this perspective but also extend it to proteins reconstituted into lipid vesicles, which appear, by an as-yet unidentified mechanism, to intrinsically favor high protein content (Fig. 1).

Importantly, under these crowded conditions, the membrane strain profile becomes highly sensitive to inter-protein spacing, and is therefore a direct function of *x*_p_. It is also noteworthy that, for the vesicle sizes considered here, membrane curvature effects are negligible at the short distances between protein inclusions, and thus do not substantially influence the observed local deformation profiles.

These detailed molecular insights provided the foundation for our SAXS modeling of OmpLA-containing vesicles. In addition to extracting 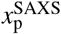, our multiscale analysis focused on quantifying the average protein-induced membrane strain, which exhibited excellent agreement with results from MD simulations. Notably, our analysis suggests that OmpLA predominantly adopts monomeric states in POPC PLUVs, consistent with the absence of Ca^2+^ ions, which are required to stabilize active OmpLA dimers,^4,21^ and with the interpretation that membrane-mediated deformations result in a weak net repulsive interaction between proteins.

For DLPC PLUVs, the relatively low effective protein concentration precluded a similar analysis. It is worth noting, however, that previous work in egg-yolk phosphatidylcholine vesicles reported mainly dimeric OmpLA states,^47^ underscoring the strong influence that lipid composition exerts on membrane-mediated protein interactions.

## VI. CONCLUSIONS

We have introduced an integrative methodology that combines all-atom MD simulations with multiscale SAXS analysis to quantify the structural response of lipid-bilayer membranes to the inclusion of an integral membrane protein. For OmpLA reconstituted into ∼100 nm POPC vesicles, we obtained protein/lipid ratios from SAXS that agree quantitatively with independent optical measurements and average membrane strain values that match MD predictions within experimental uncertainty.

MD simulations reveal that OmpLA induces oscillatory thickness deformations extending up to eight lipid shells from the protein boundary, yet the net change in average membrane thickness remains below 1%. Detecting such subtle effects by SAXS alone would not be feasible, as they fall below the typical relative uncertainty of thickness measurements using scattering.^18,32^ This challenge was addressed by employing constrained Bayesian optimization within the *SAS_MoCa* framework,^38,39^ which enables the incorporation of information from different sources, such as MD, into the data analyses.

Our approach proved less effective for DLPC-based proteoliposomes, where substantial protein loss after reconstitution limited the analysis. Overcoming this limitation will require improved sample preparation strategies or the use of lipid systems with intermediate hydrophobic thickness.

Ongoing efforts aim to extend the scattering analysis by incorporating contrast variation from small-angle neutron scattering and complementary structural constraints from nuclear magnetic resonance spectroscopy, as well as accounting for vesicle dynamics and bending fluctuations^48,49^. Together, these developments will enable quantitative investigations of protein–lipid coupling in increasingly complex biomembrane systems.

## SUPPLEMENTARY MATERIAL

## ACKNOWLEDGMENTS

This work was supported by the Austrian Science Funds (FWF) (grant-DOIs: 10.55776/P32514, and 10.55776/PIN7380924), the Austrian Research Promotion Agency (FFG) Grant 870454, the Czech Science Foundation (grant no. 25-19982L), and the project National Institute of virology and bacteriology (Programme EXCELES, ID Project No. LX22NPO5103) - Funded by the European Union - Next Generation EU. Computational resources were provided by the CESNET, CERIT Scientific Cloud, and IT4 Innovations National Supercomputing Center by MEYS CR through the e-INFRA CZ (ID:90254). POPC based samples were measured at the BioSAXS beamline P12 operated by EMBL Hamburg at the PETRA III storage ring (DESY, Hamburg, Germany), experiment R16-27, with the assistance of Haydyn Mertens. We thank Barbara Eicher and Anna Weitzer for running POPC sample measurements, and Moritz Frewein and James Jennings for the helpful discussion.

## DATA AVAILABILITY STATEMENT

SAXS data can be downloaded from https://doi.org/10.5281/zenodo.20927456. MD input and analysis files can be accessed via https://doi.org/10.5281/zenodo.21042438.

## Appendix A: Bilayer scattering amplitude

The used analytical function of the scattering amplitude of a planar lipid bilayer, *A*_M_(*q*), applies the latest update of the SDP model.^18^ The following formulas specifically refer to the parsing scheme for PC lipids,^32^ as illustrated in Figure 9.

**FIG. 9.**
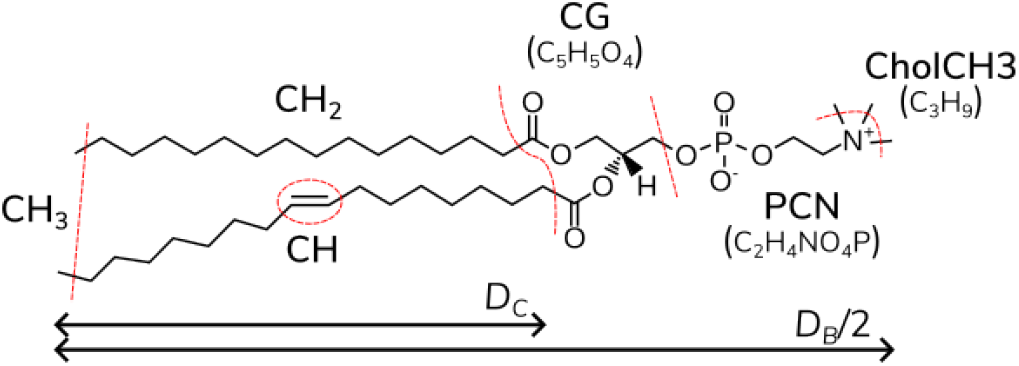
SDP parsing scheme for POPC, illustrating the defined quasimolecular groups,together with membrane structural parameters, such as the hydrocarbon chain length (*D*_C_) and half of the Luzzati thickness (*D*_B_) of the lipid bilayer (see arrows).

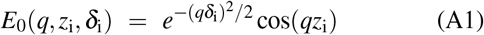

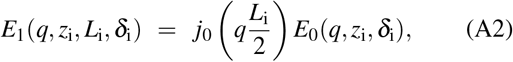

which are the Fourier transforms of a Gaussian profile (A1) and a error-function-based smooth rectangular profile (A2). Here, *z*_i_ and *δ*_i_ are, respectively, the center and the width of the Gaussian or the smooth rectangular functions, and *L*_i_ is the distance between the centers of the two error functions that comprise the smooth rectangular profile. The subscript ‘i’ refers to a given quasimolecular group.

The scattering amplitude is then given by:

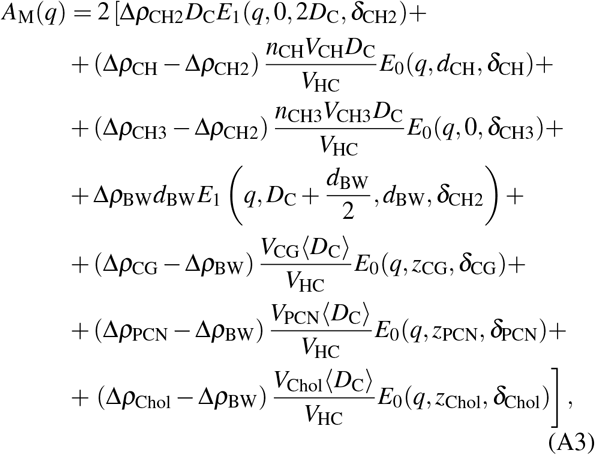

where

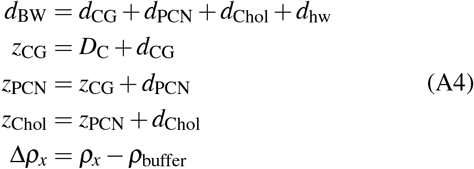

Note that the SLD of the buffer, *ρ*_buffer_, includes effects from temperature and composition (here: 20 mM TRIS and 2 mM EDTA).

Thickness fluctuations resulting from peristaltic modes are accounted by averaging the acyl-chain thickness via

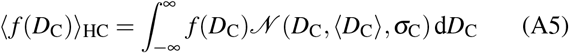

where *N* (*D*_C_, ⟨*D*_C_⟩, *σ*_C_) is the Gaussian probability distribution function of mean ⟨*D*_C_⟩ and standard deviation *σ*_C_.^18^

All parameters used for *A*_M_ are detailed in Tables II and III, see also^17,18^. The lipid volume *V*_L_ is adapted from^32^.

**TABLE I.** Estimated values for the radius of the membrane deformation zone (*r*^∗^), number of perturbed lipids 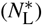, and average deformation per protein inclusion (⟨Δ⟩^∗^) for low- and high-*x*_p_ regimes (*a*_L_ ∼ 66^2^).

| System | $r^*$ (Å) | $N_L^*$ | $\langle \Delta \rangle_{\text{low-}x_p}^*$ | $\langle \Delta \rangle_{\text{high-}x_p}^*$ |
| --- | --- | --- | --- | --- |
| POPC<sup>a</sup> | 75.7 | 520 | $+0.002 \pm 0.003$ | $-0.013 \pm 0.002$ |
| DLPC<sup>a</sup> | 65.8 | 400 | $+0.002 \pm 0.003$ | $+0.007 \pm 0.005$ |
| POPC<sup>b</sup> | 87.3 | 690 | $-0.001 \pm 0.002$ | $-0.003 \pm 0.004$ |
| DLPC<sup>b</sup> | 74.2 | 510 | $+0.007 \pm 0.004$ | $+0.001 \pm 0.007$ |

<sup>a</sup> OmpLA monomer
<sup>b</sup> OmpLA dimer

**TABLE II.** List of quasi-molecular groups for PC lipids.^17,18^.

| Abbreviation | Content |
| --- | --- |
| CH3 | Terminal methyl group |
| CH2 | Methylene group(s) |
| CH | Methine group |
| CG | Carbonyl-glycerol backbone |
| PCN | Phosphate group + CN |
| Chol | Choline-CH3 group |
| BW | Hydration layer |

**TABLE III.** Recap of the structural parameters of interest relative to *A*M(*q*).^17,18^.

| Parameter | Description |
| --- | --- |
| $\rho_x$ | SLD of group $x$ |
| $d_x$ | Relative position of group $x$ |
| $\delta_x$ | Fluctuation width of group $x$ |
| $V_x$ | Molecular volume of group $x$ |
| $n_x$ | Number of group $x$ units |
| $d_{BW}$ | Width of hydration layer |
| $2D_C$ | Hydrophobic thickness |
| $\sigma_C$ | Acyl-chain thickness fluctuation width |
| $a_L = V_{HC}/D_C$ | Lateral area per lipid |
| $D_B = 2V_L/a_L$ | Luzzati bilayer thickness |
| $V_{HC} = V_L - V_H$ | Lipid acyl-chain volume |
| $V_L$ | Lipid volume |
| $V_H$ | Lipid head-group volume |
| $r_{32} = V_{CH3}/V_{CH2}$ | CH<sub>3</sub> to CH<sub>2</sub> volume ratio |
| $r_{12} = V_{CH}/V_{CH2}$ | CH to CH<sub>2</sub> volume ratio |
| $r_{CG} = V_{CG}/V_H$ | CG rel. volume |
| $r_{PCN} = V_{PCN}/V_H$ | PCN rel. volume |
| $V_{BW}$ | Hydration-layer water volume |

Note that the SLD of the buffer, $\rho_{\text{buffer}}$ , includes effects from temperature and composition (here: 20 mM TRIS and 2 mM EDTA).

## Appendix B: Size polydispersity of vesicles

The variation of the PLUV radius, *R*_v_, is described by the Schulz probability distribution function (pdf) (see, e.g., 33). The Schulz pdf enables the analytical calculation of its *n*-th moments 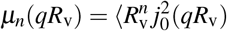:

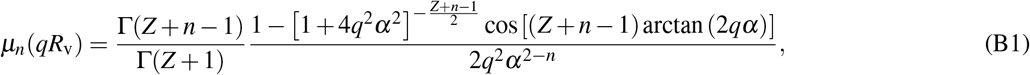

where 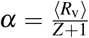, and *Z* is the width parameter of the Schulz pdf. The variance is given by *σ* ^2^ = ⟨ *R*_v_⟩ ^2^*/*(*Z* + 1) and the forward scattering limit is

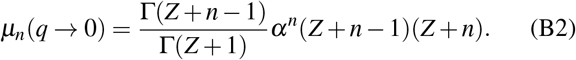

Below we detail the zeroth, second and forth moments of this pdf:

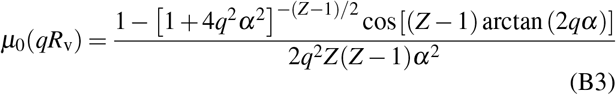

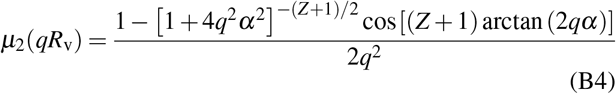

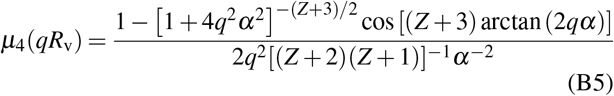

## Appendix C: Scattering amplitude of the virtual protein model

The amplitude of the virtual protein model is represented by a cylindrically symmetric stack of 2M + 1 coaxial disks that mimic an OmpLA monomer (or dimer), *A*_disks_, capped by two additional disks representing the protein hydration layers, *A*_hw_. The disks are stacked along the membrane normal, with the membrane midplane serving as a mirror plane of symmetry:

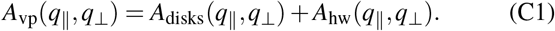

The scattering amplitude of the protein term is

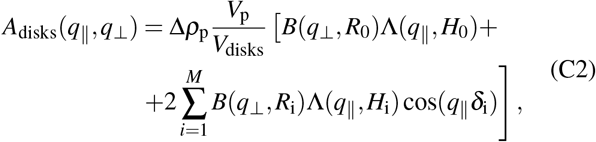

where *R*_i_ and *H*_i_ are the different radii and lengths of the disks, *R*_0_ and *H*_0_ refer to the single central disk, and

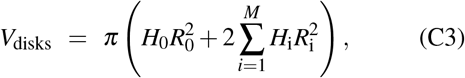

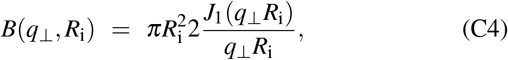

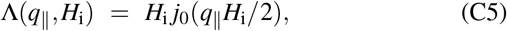

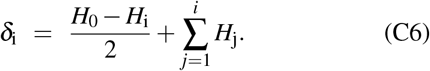

Briefly, *V*_disks_ is the total volume of all disks, which is used to re-normalize the virtual-protein to the actual volume, *V*_p_, of an OmpLA monomer (or dimer). *B* and Λ functions are the decoupled radial and axial partial scattering amplitudes, and *δ*_i_ is the center-of-mass distance between the central and the *i*^th^ disk.

Analogously, the scattering amplitude of the two hydration disks is

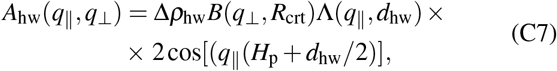

where the disk radii are estimated from the average crosssection of OmpLA monomer (or dimer), *R*_crt_, and their thicknesses correspond to one molecular layer of water, *d*_hw_. Note that their position depends on the length of OmpLA along the membrane axis, *H*_p_, and is fixed to ± (*H*_p_ + *d*_hw_*/*2). The parameters defining the virtual protein model for OmpLA monomers and dimers are listed in Tables IV and V.

**TABLE IV.** Summary of structural parameters describing OmpLA, including SLDs. Values were calculated from the PDB structures 1qd5 (monomers) and 1qd6 (dimers).^21^.

| Parameter | Value |
| --- | --- |
| $H_p$ | 44.0 Å |
| $R_{\text{crt}}(\text{monomer})$ | 17.2 Å |
| $R_{\text{crt}}(\text{dimer})$ | 20.3 Å |
| $\rho_p$ | $12.02 \times 10^{-6} \text{ Å}^{-2}$ |
| $\rho_{\text{hw}}$ | $11.50 \times 10^{-6} \text{ Å}^{-2}$ |
| $V_p(\text{monomer})$ | $3.8906 \times 10^4 \text{ Å}^3$ |
| $V_p(\text{dimer})$ | $7.1932 \times 10^4 \text{ Å}^3$ |

**TABLE V.** List of normalized parameters defining the virtual protein model of OmpLA monomers and dimers (Eq. C2). Disk radii are normalized by the OmpLA cross section, 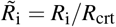, and lengths by the even fractions of OmpLA length, 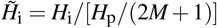.

| | Monomer, $M = 6$ | Dimer, $M = 4$ |
| --- | --- | --- |
| $(\tilde{R}_0, \tilde{H}_0) =$ | (2.0972, 0.9586) | (2.1459, 0.5392) |
| $(\tilde{R}_1, \tilde{H}_1) =$ | (1.1590, 1.3194) | (0.7683, 1.0556) |
| $(\tilde{R}_2, \tilde{H}_2) =$ | (1.0291, 1.0394) | (0.9526, 1.3101) |
| $(\tilde{R}_3, \tilde{H}_3) =$ | (0.3677, 1.1493) | (1.2479, 1.1562) |
| $(\tilde{R}_4, \tilde{H}_4) =$ | (0.4156, 1.0470) | (0.9900, 1.1711) |
| $(\tilde{R}_5, \tilde{H}_5) =$ | (0.4510, 1.2370) | - |
| $(\tilde{R}_6, \tilde{H}_6) =$ | (0.3907, 1.3432) | - |

## Appendix D: Protein-protein structure factor

The full protein term derived from Eq. 4 is:

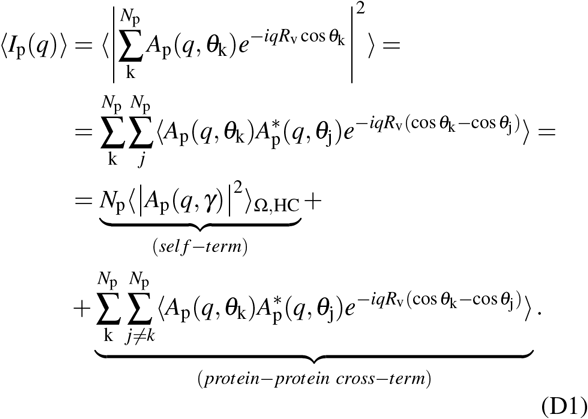

The *self-term* corresponds to the protein form factor, where the information about protein positions is lost upon averaging. Note that angular brackets explicitly not only refer to the orientational average of the membrane proteins (where *γ* is the angle between **q** and the protein director, Fig. 3), but also to averaging emerging from membrane thickness fluctuations. In contrast, the *protein-protein cross-term* does depend on protein positions and orientations. However, if we assume that the proteins are distributed homogeneously on the PLUV surface with their directors oriented parallel to the radius of the PLUV, then the cross-term will be dominated by the average distance between the proteins. This allows us to factorize the ensemble averages over the distances and orientations^36^. Thus, Equation D1 can be approximated by

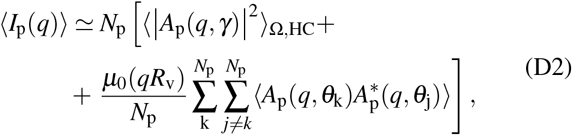

where the center-of-mass distribution of proteins contributes to the structure factor with the term *µ*_0_(*qR*_v_)^19^ (Eq. B3), and also includes PLUV polydispersity. A further approximation of the double sum in Eq. (D2) leads to the ensemble average of the protein term^36^:

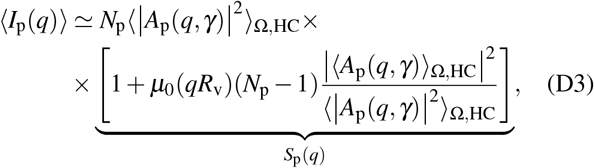

where *S*_p_(*q*) is the effective structure factor of the protein-protein term.

## Notes

### Competing Interest Statement

The authors have declared no competing interest.

