## SUPPLEMENTARY MATERIAL for "Membrane Thickness Strain from Protein Inclusions: A Multiscale Simulation and X-Ray Scattering Study of Proteoliposomes"

### SUPPLEMENTARY MATERIAL - Linking Membrane Deformations and Protein Copy Number: An Integrated Molecular Dynamics and X-Ray Scattering Approach

Enrico F. Semeraro,[1,2,3] Ladislav Bartoš,[4,5] Paulina Piller,[1] Rahul Deb,[4,5] Sandro Keller,[1,2,3] Robert Vácha,[4,5] and Georg Pabst[1,2,3,a)]

[1)] *Department of Molecular Biosciences, NAWI Graz – University of Graz, Graz, Austria*

[2)] *Field of Excellence BioHealth – University of Graz, Graz, Austria*

[3)] *BioTechMed Graz, Graz, Austria*

[4)] *CEITEC – Central European Institute of Technology – Masaryk University, Brno, Czech Republic*

[5)] *National Centre for Biomolecular Research – Masaryk University, Brno, Czech Republic*

(*Electronic mail:.)

(Dated: 3 July 2026)

[a)] https://membrane-biophysics.uni-graz.at/en/.

#### S1. MATERIALS AND SAMPLE PREPARATION

##### Chemicals

1-palmitoyl-2-oleoyl-sn-glycero-3-phosphocholine (POPC), 1,2-dilauroyl-sn-glycero-3-phosphocholine (DLPC), 1-palmitoyl-2-oleoyl-sn-glycero-3-phospho-(1'-rac-glycerol) (POPG), and 1,2-dilauroyl-sn-glycero-3-phospho-(1'-rac-glycerol) (DLPG) were purchased from Avanti Polar Lipids (Alabaster, AL, USA). Isopropyl-$\beta$-d-thiogalacto-pyranoside (IPTG), tris(hydroxymethyl) aminomethane (Tris), D(+)-saccharose, urea, glycine, KCl, $H_2SO_4$ (96 % v/v), $Na_2S_2O_5$ and $Na_2SO_3$ were obtained from Carl Roth (Karlsruhe, Germany). Lauryldimethylamin-N-oxid (LDAO), $HClO_4$ (67−72 % v/v) and ammonium heptamolybdate tetrahydrate were acquired from Sigma-Aldrich (Vienna, Austria). Ethylendiaminetetraacetatic acid (EDTA), 1-amino-2-hydroxynaphthalin-4-sulfonic acid (ANSA) and $KH_2PO_4$ from Merck (Darmstadt, Germany).

##### OmpLA Production and Purification

OmpLA without signal sequence was expressed in *Escherichia coli* BL21(DE3) cells (Novagen, Merck, Darmstadt, Germany) as inclusion bodies (IBs) utilizing a pET-24a (+) expression vector (Merck, Darmstadt, Germany). After inoculation of the lysogeny broth medium with *E. coli* BL21(DE3) cells, the production of OmpLA was induced by the addition of IPTG (0.4 mM final concentration), followed by an incubation period of 3 h at 37 °C under agitation. Harvested cells were washed and resuspended in 1:10 (wt/vol) ice-cold breakage buffer (50 mM Tris, 40 mM EDTA, 25% (wt/vol) saccharose, pH 8.0). Then, cells were sonicated with a sonopuls HD 2070 homogenizer (Bandelin, Berlin, Germany) for 10 min (pulse: 30 s, pause: 30 s) at an amplitude of 40%. The cell lysate was centrifuged for 1 h at 4 °C and 7000 g, the pellet was resuspended in washing buffer (10 mM Tris, 1 mM EDTA, pH 8.0), centrifuged again with the same settings and dissolved overnight in solubilization buffer (20 mM Tris, 2 mM EDTA, 8 M urea, 100 mM glycine) at 4 °C under agitation. Solubilized IBs were centrifuged for 30 min at 4 °C and 7000 g. The protein concentration of the supernatant was determined with a Nanodrop ND-1000 spectrophotometer (Peqlab Biotechnology GmbH, Erlangen, Germany) by measuring the absorbance ($A$) at 280 nm, and using the reference value at 10 g/l, $A^{1\%}_{\mathrm{OmpLA}} = 26.68$ (calculated from ProtParam, Expasy, the Swiss Bioinformatics Resource Portal). For blank measurement buffer was used.

Refolding of OmpLA was executed by adding the refolding buffer by drop dilution under agitation at 50 °C to reach final concentrations of 0.33 mg/mL OmpLA, 20 mM Tris, 2 mM EDTA, 0.80 M urea, 10 mM glycine and 35 mM LDAO (pH 8.3). The mixture was further agitated for 16 h at 50 °C. The refolded fraction was centrifuged for 15 min at 4 °C and 7000 g and filtered through a 450-nm poly (-vinylidene fluoride) filter (Carl Roth, Karlsruhe, Germany). The folded protein fraction was separated from the unfolded one by anion exchange chromatography on a 20 ml (4x5 ml) tandem HiTrap DEAE column (GE Healthcare, Solingen, Germany) in 20 mM Tris, 2 mM EDTA, 35 mM LDAO (pH 9.5) with a two-step gradient of 105 mM and 1.5 M KCl. The pooled folded fractions were dialyzed [20 mM Tris, 2 mM EDTA, 12 mM LDAO (pH 8.3)] overnight and concentrated on a 5 ml Resource Q column (GE Healthcare, Solingen, Germany) in 20 mM Tris, 2 mM EDTA and 12 mM LDAO (pH 8.3). The protein was eluted with 20 mM Tris, 2 mM EDTA, 12 mM LDAO and 1.5 mM KCl (pH 8.3) and further desalted on a PD-10 column (GE Healthcare, Solingen, Germany) using 20 mM Tris, 2 mM EDTA and 2 mM LDAO (pH 8.3). The final protein concentration was measured as described above.

**Large Unilamellar Vesicles Preparation**

Lipids (POPC, POPE, POPG) were dispersed in a 2:1 vol/vol chloroform/methanol mixture, dried under a stream of nitrogen and stored in vacuum overnight to ensure complete solvent evaporation. For lipid hydration the reconstitution buffer (20 mM Tris, 2 mM EDTA, pH 8.3) was added. The formation of lipid vesicles was achieved by intermittent vigorous vortexing at 15 °C $> T_m$ (lipid melting temperature) for 1 h. Large Unilamellar Vesicles (LUVs) were prepared by 31-fold extrusion (mini extruder: Avanti Polar Lipids, Alabaster, AL, USA) through a polycarbonate filter (Whatman Nuclepore™ Track-Etched Membranes from Merck, Darmstadt, Germany) with a pore diameter of 100 nm. LUV formation was assisted by doping the bilayers with 5 mol% POPG[?] . POPG does not affect membrane structure or protein activity at presently low concentrations (see below). Vesicle size was checked with dynamic light scattering (DLS) using a Zetasizer Nano ZSP (Malvern Panalytical, Malvern, UK). We found unimodal size distributions with intensity peak centered in the range $150 - 160$ *nm* (data not shown).

**Proteoliposome preparations - Protein Reconstitution**

For reconstitution, OmpLA solubilized in 2 mM LDAO was added drop by drop to the lipid vesicles at 35 °C and 350 rpm in a thermomixer (Eppendorf, Hamburg, Germany) to reach a final lipid/protein mole ratio of 300:1 (600:1 and 900:1, respectively). LDAO was removed by dialysis against 50 mM Tris, 2 mM EDTA and 200 mM NaCl (pH 8.3) overnight. The proteoliposomes were extruded 31 times through a 100 nm polycarbonate filter. Similarly to LUVs, we found of PLUVs unimodal size distributions in the range at $z_{av} = 160 - 170\ nm$ (data not shown).

**Phosphate Assay**

The lipid samples were carbonized first in a heated metal block at maximum heat. After cooling, 0.4 ml acid mixture (9:1 vol/vol conc. $H_2SO_4$:$HClO_4$) was added and the sample was heated up again for 30 min. 9.6 ml of a reaction mixture (22 ml reagent A: 10.5 mM ANSA, 0.7 mM $Na_2S_2O_5$, 40 mM $Na_2SO_3$; 500 ml reagent B: ammonium heptamolybdate tetrahydrate (0.26 %)) was added to the cool samples. After mixing, the tubes were placed in a sand bath at 90 °C for 20 min. The extinction of the cooled samples was measured on a Spectrophotometer Onda V-10 Plus (Labbox Labware, S.L., Barcelona, Spain) at 830 nm. The phospholipid concentration was calculated by using a calibration curve (3.2 mM $KH_2PO_4$ as phosphate-standard-solution containing $1 - 14\ \mu$g phosphor).

#### S2. MOLECULAR DYNAMICS SIMULATIONS

*a. General information* All molecular dynamics simulations for quantifying membrane deformation were performed using the GROMACS simulation package version 2021.4[1] with the CHARMM36m force field[2,3] describing lipids and proteins and with the TIP3P model[4] describing water. The structures of the OmpLA dimer and monomer solved by X-ray diffraction were obtained from the RCSB Protein Data Bank, entries 1QD6 (www.rcsb.org/structure/1QD6) and 1QD5 (www.rcsb.org/structure/1QD5), respectively. Missing N-terminal loop segments were reconstructed using MODELLER version 10.4[5]. The entire protein structure was then minimized in vacuum using the steepest-descent algorithm until the maximum force decreased below 100 kJ $mol^{-1}$ $nm^{-1}$.

*b. Systems preparation* Using CHARMM-GUI[6,7], we prepared several membrane systems composed of POPC (1-palmitoyl-2-oleoyl-*sn*-glycero-3-phosphocholine) or DLPC (1,2-dilauroyl-*sn*-glycero-3-phosphocholine) lipids with embedded monomeric or dimeric OmpLA. For each lipid type and for both monomeric and dimeric OmpLA, two types of membrane systems were constructed. To generate the *low-$x_p$ regime*, a single protein unit was embedded in a large membrane containing 800 lipid molecules for monomers and 1200 for dimers. Conversely, to generate the *high-$x_p$ regime*, a single protein unit was embedded in a smaller membrane composed of 200 lipids for monomers and 500 for dimers. Each system also contained NaCl ions at a physiological concentration (0.154 mol $dm^{-3}$) with an excess of ions to neutralize the system.

*c. Minimization* After system construction, we minimized each system using the steepest-descent algorithm with a force tolerance of 1000 kJ $mol^{-1}$ $nm^{-1}$. During the minimization, positional restraints were applied to the protein backbone and protein side chains (force constants of 4000 and 2000 kJ $mol^{-1}$ $nm^{-2}$, respectively) as well as to the phosphorus atom of each lipid (force constant of 1000 kJ $mol^{-1}$ $nm^{-2}$). Additional restraint (force constant of 1000 kJ $mol^{-1}$ $rad^{-2}$) was applied to the dihedral between C1, C3, C2, and O21 of all lipid molecules (glycerol carbons and oxygen linking the *sn*-2 tail to glycerol), fixing it at $-120 \pm 2.5°$. In systems containing POPC, an additional dihedral restraint of the same strength was applied to the dihedral between C28, C29, C210, and C211 (carbons around the double bond of the oleoyl tail), fixing it at $0 \pm 0.0°$.

*d. Equilibration* Equilibration was performed in six stages of different simulation lengths: I-III) 250 ps each, IV-V) 1 ns each, VI) 5 ns. In stages I-III, the simulation time step was 1 fs, while in the rest of the equilibration (and the following production stage), the time step was 2 fs. Stages I and II were performed in the canonical (NVT) ensemble, and the other stages in the isothermal-isobaric (NPT) ensemble. The equations of motion were integrated using the leap-frog algorithm. A stochastic velocity rescaling thermostat[8] with a coupling constant of 0.5 ps was employed to maintain the temperature at 310 K in three separate thermal baths (protein, membrane, and water with ions). In the NPT stages of equilibration, the pressure was kept at 1 bar using the Berendsen barostat[9] with a semi-isotropic coupling scheme to independently scale the simulation box in the *xy*-plane (the membrane plane) and along the *z*-axis, a coupling constant of 5 ps, and a compressibility of $4.5\times10^{-5}$ bar$^{-1}$. Short-ranged non-bonded interactions were truncated at 1.2 nm, with a force switch applied starting from 1.0 nm. Electrostatic interactions were treated using the Fast Smooth Particle-Mesh Ewald method[10]. Bonds involving hydrogens were constrained using the LINCS algorithm[11]. Translational velocity removal was applied separately for the membrane with protein and for water with ions. The same position and dihedral restraints as in minimization were applied in equilibration stages as well, with the strength of the restraints gradually decreasing (see Table S1).

TABLE S1. Position and dihedral restraints applied during the individual stages of equilibration. All values are in kJ mol$^{-1}$ nm$^{-2}$ or kJ mol$^{-1}$ rad$^{-2}$ (for lipid dihedrals).

| | protein backbone | protein side chains | lipid phosphorus | lipid dihedrals |
|---|---|---|---|---|
| I | 4000 | 2000 | 1000 | 1000 |
| II | 2000 | 1000 | 400 | 400 |
| III | 1000 | 500 | 400 | 200 |
| IV | 500 | 200 | 200 | 200 |
| V | 200 | 50 | 40 | 100 |
| VI | 50 | 0 | 0 | 0 |

*e. Production* After equilibration, each system was simulated in three replicas with the same simulation parameters as in stage VI of equilibration, except that the Berendsen barostat was replaced with the Parrinello-Rahman barostat[12,13] and no restraints were applied. The length of the production phase for each replica was 2 $\mu$s, with the first 500 ns excluded from the analysis.

*f. Analysis* Membrane thickness was calculated using an in-house developed tool `memthick` (available at github.com/Ladme/memdian). For all systems, we first generated two-dimensional thickness maps for each replica by creating a mesh for each membrane leaflet with $0.1\times0.1$ nm$^2$ bins and computing the average z-position of phosphate beads in each bin from the simulation trajectory. The membrane thickness for each bin was then determined as the difference between the average z-positions of phosphates in the corresponding upper- and lower-leaflet bins.

For systems in *low-*$x_p$ *regime*, we then performed radial averaging of the two-dimensional maps to obtain the membrane thickness as a function of distance from the protein's center of geometry. The one-dimensional profiles show membrane thickness *around* an independent OmpLA protein. The resulting one-dimensional profiles from individual replicas were averaged, with the standard deviation reported as the error estimate.

For systems in *high-*$x_p$ *regime*, we first expanded the system to include neighboring copies of the central simulation box. We then calculated one-dimensional membrane thickness profiles (capturing membrane thickness *between* two neighboring OmpLA proteins) by averaging the membrane thickness along the lines connecting the central protein copy with its four nearest virtual neighbors (one in each direction). The resulting one-dimensional profiles from individual replicas were averaged, with the standard deviation reported as the error estimate.

#### S3. MD - SUPPLEMENTARY FIGURES

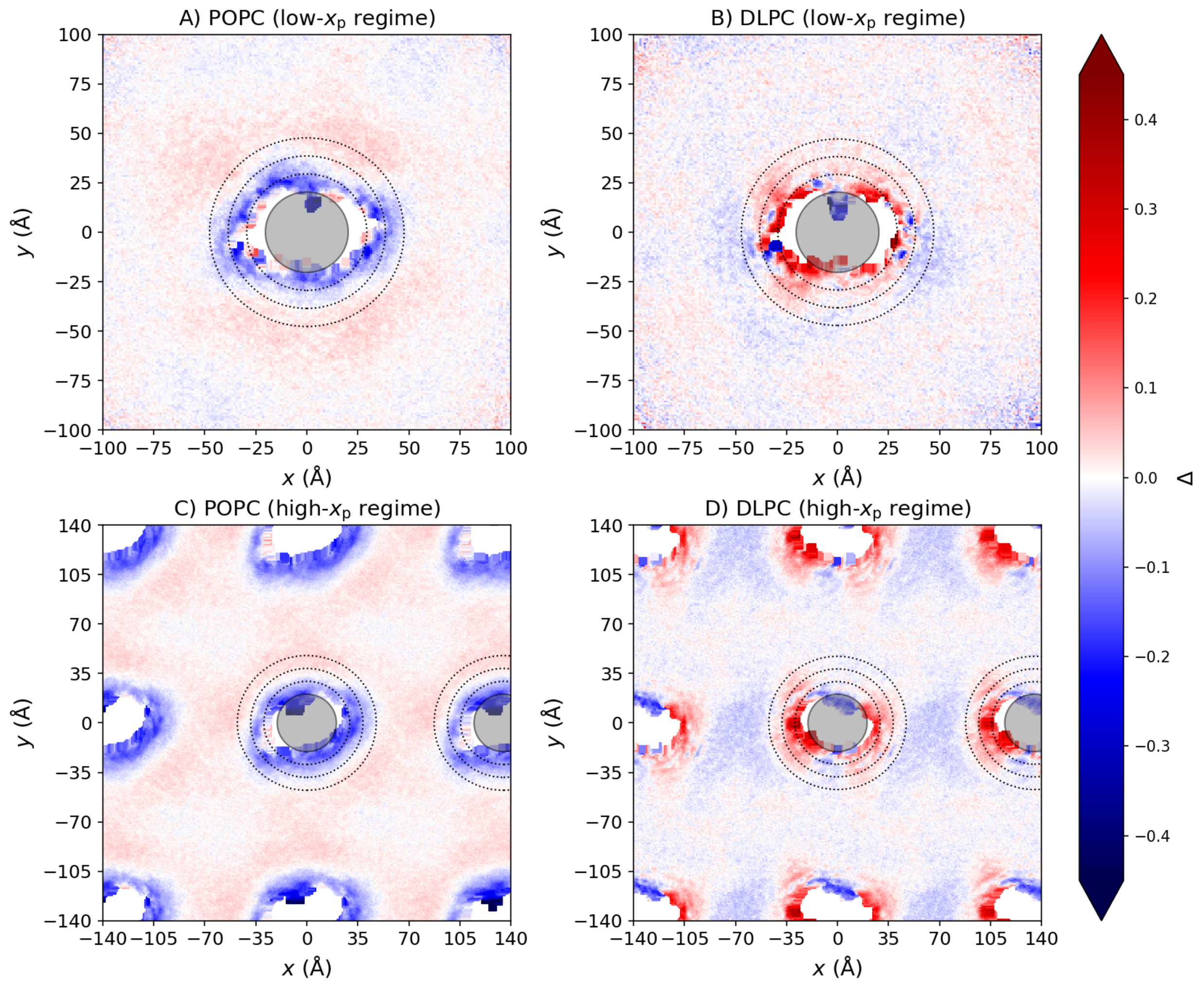


FIG. S1. Relative lipid thickness deformation Δ around an OmpLA-dimer inclusion (empty space in the center). Panels **A-B** show isolated OmpLA dimers in the low-$x_p$ regime, while panel **C-D** includes neighboring simulation boxes, simulating the crowded high-$x_p$ system. The color code highlights thickening (red) and thinning (blue). The shaded circle represents the dimer's cross section with radius $R_{crt}$, and the dotted lines mark the average boundaries of the first three lipid layers.

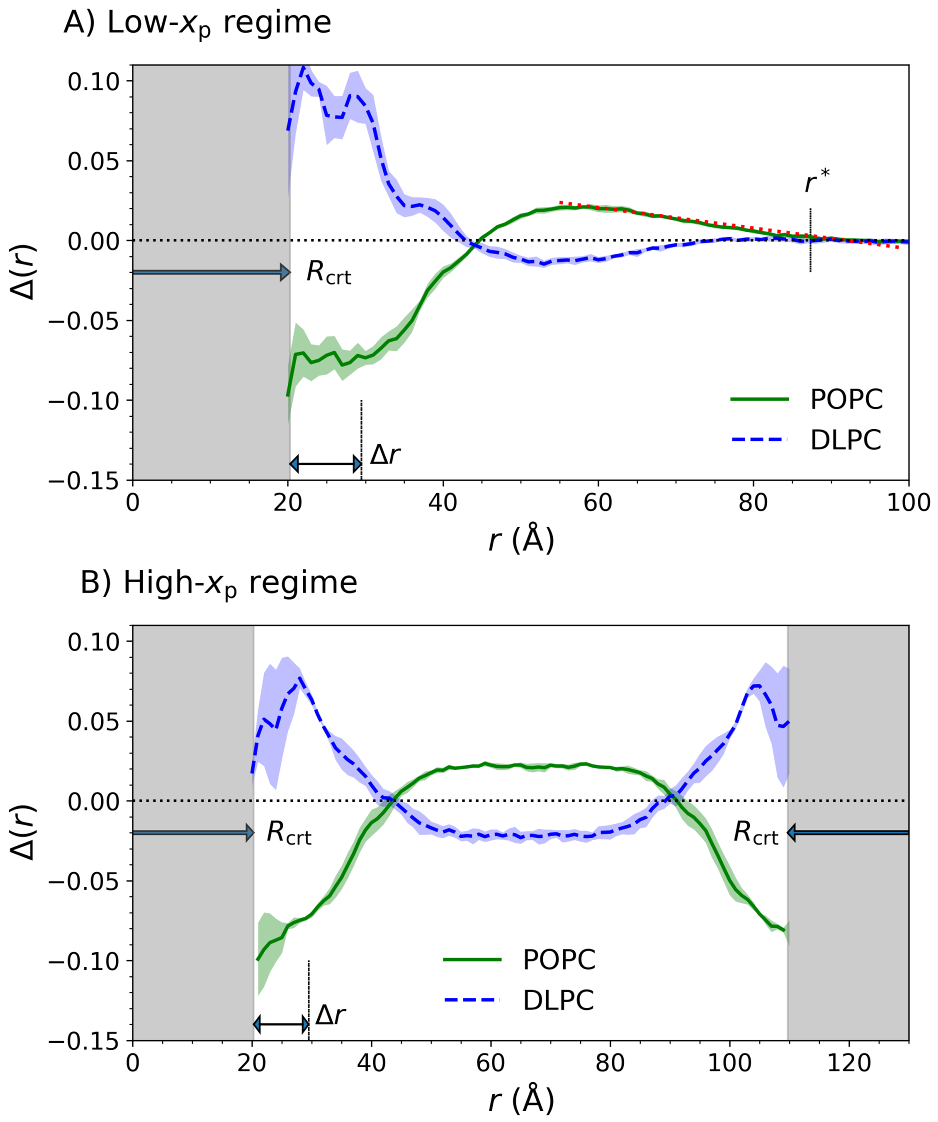


FIG. S2. Average membrane-deformation $\Delta$ profiles for POPC (solid green) and DLPC (dashed blue) as a function of distance from the OmpLA dimer centre of mass (shaded bands indicate the associated standard-error envelopes). Each curve represents the average over three independent replicas; $\Delta = 0$ corresponds to bulk lipid. The gray shaded region marks the spatial extent of a single OmpLA dimer, whose mean radius is $R_{crt} = 20.3$ Å. $\Delta r$ is the average diameter of a lipid and illustrate the extension of the annular layer. **(A)** Low-$x_p$ regime: radial averaging around the OmpLA inclusion. The dotted red line denotes the zero-crossing used to determine the maximal radial reach of the deformation, $r^*$ (illustrated for POPC only). **(B)** High-$x_p$ regime: linear averaging performed along the line connecting two OmpLA-dimer inclusions.

#### S4. SAXS ANALYSIS - CURVE FITTING INITIALIZATIONS

SAXS curves were analyzed with the *SAS_MoCa* tool ([14,15]). For each adjustable parameter, $y$, one must initialize either (i) a non-informative prior, that is a range of possible values in the range $[y_{min}, y_{max}]$, or (ii) an informative prior, that is a Gaussian profile $\mathcal{N}(\bar{y}, \sigma_{\bar{y}}^2)$ with mean $\bar{y}$ and standard deviation $\sigma_{\bar{y}}$.

Most of the priors for the SDP parameters of all POPC and DLPC curves (mock and pLUVs) were selected by combining published results ([16,17]) and interpolate them at 37 °C. The only exception is the lipid head-group volume, $V_{\mathrm{H}}$, whose prior describes the current precision in its determination, according to[18]. These are summarized in Table S2.

In contrast, the remaining adjustable parameters required tailored informative or non-informative priors initializations, that are summarized in Tables S3, and S4. The priors' details for $x_{\mathrm{p}}^{\mathrm{SAXS}}$ parameters are inherited from the experimental determination of OmpLA-to-lipid molar fraction, as described in the manuscript. For the other relevant parameters, the prior choices are summarized in the following.

##### A. Membrane thickness

We used the reference bulk values, $2D_{\mathrm{C}}^{0}$, and associated error from[16,17] to set the Gaussian prior for both POPC and DLPC mock LUVs (values for 37 °C were obtained via interpolation). These are $\bar{y} = 28.6$ Å for POPC, and $\bar{y} = 21.4$ Å for DLPC, both with a relative standard deviation of 3%.

For pLUV systems instead, we used the membrane thickness result obtained from our reference samples, $2D_{\mathrm{C}}^{0,mock}$ (mock LUVs), and leveraged the predicted $\langle\Delta\rangle$ values obtained from MD simulations. That is, for the mean of the normal prior we used $\bar{y} = 2D_{\mathrm{C}}^{0,mock}(\langle\Delta\rangle + 1)$. The priors' $\sigma_{\bar{y}}$ for pLUVs were set to an error value of 3%.

##### B. Membrane thickness fluctuation

We applied a Gaussian prior with $\bar{y} = 0.079$ and relative $\sigma_{\bar{y}}$ of 6% for the relative thickness fluctuation, $\varepsilon_{\mathrm{C}}$, of mock LUVs data, based on the results reported in[17]. However, we assumed that the membrane dynamics in pLUVs, particularly in the high-$x_{\mathrm{p}}$ regime, may differ from those in LUVs. In this case, therefore, we used a non-informative prior with boundaries $[0.055, 0.2]$.

#### C. Hydration layer

For the mock LUV data, we assigned a non-informative prior with boundaries $[28,32]\ ^3$ to the volume of water molecules in the hydration layer of the lipid membrane, $V_{BW}$.

However, for pLUV systems, we designed a prior scheme that includes the hydration shell of OmpLA, as the water molecules in the lipid-head group at the interface with OmpLA are likely to be affected by the neighboring amino-acid residues. Specifically, we assigned to $\bar{y}$ the weighted average

$$\bar{y} = V_{\mathrm{BW}}^{0}(1 - N_{\mathrm{L}}^{\mathrm{ann}} x_{\mathrm{p}}^{\mathrm{SAXS}}) + V_{\mathrm{hw}} N_{\mathrm{L}}^{\mathrm{ann}} x_{\mathrm{p}}^{\mathrm{SAXS}}, \tag{1}$$

where $V_{\mathrm{BW}}^{0} = 29.9\ ^3$ according to[17], $V_{\mathrm{hw}} = 24.5\ ^3$ is the volume of water molecules in the protein hydration layer[19], and $N_{\mathrm{L}}^{\mathrm{ann}}$ is the number of annular lipids:

$$N_{\mathrm{L}}^{\mathrm{ann}} = \frac{2\pi}{a_{\mathrm{L}}} \int_{R_{\mathrm{crt}}}^{R_{\mathrm{crt}}+\Delta r} r\,\mathrm{d}r. \tag{2}$$

where we used $a_{\mathrm{L}} \sim 66$ and $\Delta r \sim 9$ as first approximation values in the case of both POPC and DLPC. Note that the prior center hence depends on $x_{\mathrm{p}}^{\mathrm{SAXS}}$ and, most importantly, it represents a maximum estimate. In other words, we assign the same water-volume value to the annular lipid region as we would to an isolated protein. Given the low relative error obtained for $V_{BW}$ parameters, we assigned to the priors an arbitrary small relative $\sigma_{\bar{y}}$ of 1%.

#### S5. SAXS ANALYSIS - PRIOR SCHEMES

TABLE S2. Set of values for the shared normal priors, $\mathcal{N}(\bar{y}, \sigma_{\bar{y}}^2)$, and fixed values for POPC and DLPC systems. The table shows the prior centers, $\bar{y}$ ,and its relative standard deviation, $\sigma_{\bar{y}}/\bar{y}$, in parentheses.

| **Parameter** $y$ | **POPC** | **DLPC** |
|---|---|---|
| $d_{\mathrm{cholCH3}}$ (Å) | 2.7 (2%) | 1.8 (2%) |
| $\sigma_{\mathrm{cholCH3}}$ (Å) | 3.0 (2%) | 3.0 (2%) |
| $d_{\mathrm{PCN}}$ (Å) | 4.1 (8%) | 4.0 (8%) |
| $\sigma_{\mathrm{PCN}}$ (Å) | 2.5 (20%) | 2.5 (20%) |
| $d_{\mathrm{CG}}$ (Å) | 0.8 (8%) | 1.0 (8%) |
| $\sigma_{\mathrm{CG}}$ (Å) | 2.5 (20%) | 2.5 (20%) |
| $d_{\mathrm{CH}}$ (Å) | 9.0[a] | - |
| $\sigma_{\mathrm{CH}}$ (Å) | 3.05[a] | - |
| $\sigma_{\mathrm{CH2}}$ (Å) | 2.5 (2%) | 2.5 (2%) |
| $\sigma_{\mathrm{CH3}}$ (Å) | 3.4 (5%) | 1.54 (5%) |
| $r_{\mathrm{PCN}}$ | 0.29 (2%) | 0.26 (2%) |
| $r_{\mathrm{CG}}$ | 0.45 (2%) | 0.48 (2%) |
| $r_{12}$ | 0.81[†] | - |
| $r_{32}$ | 2.09 (2%) | 1.93 (2%) |
| $V_{\mathrm{H}}$ (Å$^3$) | 324 (2%) | 324 (2%) |

[a] Fixed values

TABLE S3. POPC systems - Set of tailored priors. The mean value of the $x_{\mathrm{p}}^{\mathrm{SAXS}}$-prior is halved for OmpLA dimers.

| | | PLUV | PLUV | PLUV |
|---|---|---|---|---|
| **Parameter** $y$ | LUV mock | $\hat{x}_{\mathrm{p}} = 1/900$ | $\hat{x}_{\mathrm{p}} = 1/600$ | $\hat{x}_{\mathrm{p}} = 1/300$ |
| $R_{\mathrm{v}}$ (Å) | [140, 260] | [290, 490] | [290, 490] | [290, 490] |
| $Z$ | [5.1, 18] | 5.1[a] | 5.1[a] | 5.1[a] |
| $x_{\mathrm{p}}^{\mathrm{SAXS}}$ | - | 0.0038 (20%) | 0.0045 (20%) | 0.0061 (20%) |
| $2D_{\mathrm{C}}$ (Å) | 28.6 (3%) | 27.5 (3%) | 27.5 (3%) | 27.5 (3%) |
| $\varepsilon_{\mathrm{C}}$ | 0.079 (6%) | [0.055, 0.2] | [0.055, 0.2] | [0.055, 0.2] |
| $V_{\mathrm{BW}}$ (Å$^3$) | [28, 32] | 29.1 (1%) | 29.0 (1%) | 28.8 (1%) |

[a] Fixed values

TABLE S4. DLPC systems - Set of tailored priors. The mean value of the $x_{\mathrm{p}}^{\mathrm{SAXS}}$-prior is halved for OmpLA dimers.

| **Parameter** $y$ | LUV mock | PLUV $\hat{x}_{\mathrm{p}} = 1/1200$ | PLUV $\hat{x}_{\mathrm{p}} = 1/300$ |
|---|---|---|---|
| $R_{\mathrm{v}}$ (Å) | $[260, 340]$ | $[290, 450]$ | $[360, 500]$ |
| $Z$ | $[8, 30]$ | $[5.1, 15]$ | $[8, 30]$ |
| $x_{\mathrm{p}}^{\mathrm{SAXS}}$ | - | 0.0033 (30%) | 0.0061 (20%) |
| $2D_{\mathrm{C}}$ (Å) | 21.4 (2%) | 20.8 (2%) | 20.8 (2%) |
| $\varepsilon_{\mathrm{C}}$ | 0.079 (6%) | $[0.04, 0.16]$ | $[0.04, 0.16]$ |
| $V_{\mathrm{BW}}$ (Å$^3$) | $[28, 32]$ | 29.8 (1%) | 29.7 (1%) |

#### S6. SAXS ANALYSIS - RESULTS

TABLE S5. **POPC** systems - Summary of fit results for LUV mock system and pLUVs with OmpLA **monomers**. Number of iterations $N_I = 100$.

| Parameter | LUV mock | PLUV $\hat{x}_p = 1/900$ | PLUV $\hat{x}_p = 1/600$ | PLUV $\hat{x}_p = 1/300$ |
|---|---|---|---|---|
| $R_v$ (Å) | 213 (8%) | 393 (5%) | 381 (5%) | 380 (5%) |
| $Z$ | 10 (30%) | 5.1[a] | 5.1[a] | 5.1[a] |
| $x_p^{SAXS}$ | - | 0.00394 (5.0%) | 0.0046 (4.0%) | 0.0055 (5.0%) |
| $d_{cholCH3}$ (Å) | 2.7 (5%) | 2.70 (2%) | 2.71 (3%) | 2.70 (3%) |
| $\delta_{cholCH3}$ (Å) | 3.00 (3%) | 3.01 (3%) | 3.00 (3%) | 3.00 (2%) |
| $d_{PCN}$ (Å) | 4.1 (10%) | 4.2 (7%) | 4.2 (7%) | 4.2 (7%) |
| $\delta_{PCN}$ (Å) | 2.9 (20%) | 2.4 (20%) | 2.3 (20%) | 2.3 (20%) |
| $d_{CG}$ (Å) | 0.82 (11%) | 0.81 (9%) | 0.81 (9%) | 0.82 (9%) |
| $\delta_{CG}$ (Å) | 2.1 (10%) | 2.2 (14%) | 2.1 (10%) | 2.2 (14%) |
| $2D_C$ (Å) | 27.8 (3%) | 27.4 (2%) | 27.5 (1.8%) | 27.6 (2%) |
| $\varepsilon_C$ | 0.08 (4%) | 0.141 (12%) | 0.140 (12%) | 0.143 (11%) |
| $d_{CH}$ (Å) | 9.0[a] | 9.0[a] | 9.0[a] | 9.0[a] |
| $\delta_{CH}$ (Å) | 3.05[a] | 3.05[a] | 3.05[a] | 3.05[a] |
| $\delta_{CH2}$ (Å) | 2.52 (3%) | 2.51 (2%) | 2.50 (2%) | 2.50 (2%) |
| $\delta_{CH3}$ (Å) | 3.4 (9%) | 3.42 (6%) | 3.38 (6%) | 3.40 (5%) |
| $r_{PCN}$ | 0.291 (3%) | 0.290 (2%) | 0.291 (3%) | 0.291 (2%) |
| $r_{CG}$ | 0.448 (3%) | 0.449 (2%) | 0.449 (2%) | 0.449 (2%) |
| $r_{12}$ | 0.81[a] | 0.81[a] | 0.81[a] | 0.81[a] |
| $r_{32}$ | 2.1 (3%) | 2.09 (2%) | 2.10 (2%) | 2.09 (2%) |
| $V_H$ (Å$^3$) | 322 (3%) | 322 (2%) | 322 (3%) | 323 (2%) |
| $V_{BW}$( Å$^3$) | 29.86 (0.3%) | 28.76 (0.6%) | 28.64 (0.7%) | 28.51 (0.7%) |
| $a_L$[b] (Å$^2$) | 67.6 (3%) | 68.7 (2%) | 68.3 (2%) | 68.0 (2%) |
| $D_B$[b] (Å) | 37.3 (3%) | 36.7 (2%) | 36.9 (2%) | 37.1 (2%) |
| $D_{pp}$[b] (Å) | 35.4 (3%) | 36.5 (2%) | 36.7 (2%) | 36.9 (2%) |
| $n_w$ [b] | $\sim 13$ | $\sim 14$ | $\sim 14$ | $\sim 14$ |
| $\langle\Delta\rangle^{SAXS,b}$ | - | $-0.017$ (> 100%) | $-0.018$ (> 100%) | $-0.015$ (> 100%) |
| $N_p$[b] | - | 230 (15%) | 260 (18%) | 300 (13%) |
| $r_p$[b] (Å) | - | 103 (10%) | 96 (12%) | 89 (9%) |

[a] Fixed values

[b] Calculated values

TABLE S6. **DLPC** systems - Summary of fit results for LUV mock system and pLUVs with OmpLA **monomers**. Number of iterations $N_I = 100$.

| **Parameter** | LUV mock | PLUV $\hat{x}_p = 1/1200$ | PLUV $\hat{x}_p = 1/300$ |
|---|---|---|---|
| $R_v$ (Å) | 308 (4%) | 370 (8%) | 420 (5%) |
| $Z$ | 19 (20%) | 6.8 (20%) | 20 (19%) |
| $x_p^{SAXS}$ | - | 0.00041 (40%) | 0.0013 (15%) |
| $d_{cholCH3}$ (Å) | 1.8 (3%) | 1.8 (3%) | 1.8 (3%) |
| $\delta_{cholCH3}$ (Å) | 3.00 (4%) | 2.98 (3%) | 3.00 (3%) |
| $d_{PCN}$ (Å) | 3.5 (9%) | 3.8 (8%) | 4.0 (8%) |
| $\delta_{PCN}$ (Å) | 2.1 (19%) | 2.5 (16) | 2.7 (20%) |
| $d_{CG}$ (Å) | 1.00 (11%) | 1.00 (11%) | 1.04 (11%) |
| $\delta_{CG}$ (Å) | 2.2 (9%) | 2.2 (9%) | 2.1 (14%) |
| $2D_C$ (Å) | 20.7 (2%) | 20.3 (2%) | 20.4 (2%) |
| $\varepsilon_C$ | 0.08 (9%) | 0.078 (20%) | 0.09 (30%) |
| $\delta_{CH2}$ (Å) | 2.51 (3%) | 2.50 (2%) | 2.502 (3%) |
| $\delta_{CH3}$ (Å) | 1.53 (8%) | 1.56 (6%) | 1.55 (7%) |
| $r_{PCN}$ | 0.259 (3%) | 0.259 (2%) | 0.258 (3%) |
| $r_{CG}$ | 0.473 (3%) | 0.477 (3%) | 0.482 (4%) |
| $r_{32}$ | 1.93 (3%) | 1.93 (3%) | 1.93 (4%) |
| $V_H$ (Å$^3$) | 324 (2%) | 327 (2%) | 317 (3%) |
| $V_{BW}$( Å$^3$) | 30.20 (0.6%) | 29.80 (0.7%) | 30.71 (0.6%) |
| $a_L$[b] (Å$^2$) | 64.7 (2%) | 65.9 (2%) | 66.4 (2%) |
| $D_B$[b] (Å) | 30.7 (3%) | 30.1 (2%) | 29.9 (2%) |
| $D_{pp}$[b] (Å) | 29.1 (3%) | 28.9 (3%) | 29.4 (2%) |
| $n_W$[b] | $\sim 8$ | $\sim 9$ | $\sim 10$ |
| $\langle\Delta\rangle^{SAXS,b}$ | - | $-0.014$ ($> 100\%$) | $-0.021$ ($> 100\%$) |
| $N_p$[b] | - | 12 (70%) | 60 (20%) |
| $r_p$[b] (Å) | - | 410 (40%) | 220 (13%) |

[b] Calculated values

TABLE S7. **POPC** systems - Summary of fit results for pLUVs with OmpLA **dimers**. Number of iterations $N_I = 100$.

| **Parameter** | LUV mock | PLUV $\hat{x}_p = 1/900$ | PLUV $\hat{x}_p = 1/600$ | PLUV $\hat{x}_p = 1/300$ |
|---|---|---|---|---|
| $R_v$ (Å) | 213 (8%) | 380 (8%) | 370 (8%) | 375 (5%) |
| $Z$ | 10.0 (30%) | 5.1[a] | 5.1[a] | 5.1[a] |
| $x_p^{SAXS}$ | - | 0.00187 (8%) | 0.00218 (6%) | 0.00279 (5%) |
| $d_{cholCH3}$ (Å) | 2.70 (5%) | 2.70 (3%) | 2.71 (3%) | 2.71 (2%) |
| $\delta_{cholCH3}$ (Å) | 3.00 (3%) | 3.00 (2%) | 2.99 (3%) | 3.00 (2%) |
| $d_{PCN}$ (Å) | 4.1 (10%) | 4.2 (7%) | 4.4 (7%) | 4.2 (10%) |
| $\delta_{PCN}$ (Å) | 2.9 (20%) | 2.0 (20%) | 1.9 (30%) | 2.2 (30%) |
| $d_{CG}$ (Å) | 0.82 (11%) | 0.82 (9%) | 0.82 (11%) | 0.83 (8%) |
| $\delta_{CG}$ (Å) | 2.1 (10%) | 2.1 (14%) | 2.0 (15%) | 2.1 (14%) |
| $2D_C$ (Å) | 27.8 (3%) | 27.2 (2%) | 27.0 (2%) | 27.8 (3%) |
| $\varepsilon_C$ | 0.08 (4%) | 0.130 (15%) | 0.120 (17%) | 0.137 (11%) |
| $d_{CH}$ (Å) | 9.0[a] | 9.0[a] | 9.0[a] | 9.0[a] |
| $\delta_{CH}$ (Å) | 3.05[a] | 3.05[a] | 3.05[a] | 3.05[a] |
| $\delta_{CH2}$ (Å) | 2.52 (3%) | 2.51 (2%) | 2.50 (3%) | 2.51 (2%) |
| $\delta_{CH3}$ (Å) | 3.4 (9%) | 3.30 (6%) | 3.40 (6%) | 3.36 (6%) |
| $r_{PCN}$ | 0.291 (3%) | 0.290 (3%) | 0.290 (3%) | 0.290 (3%) |
| $r_{CG}$ | 0.448 (3%) | 0.444 (3%) | 0.446 (3%) | 0.446 (2%) |
| $r_{12}$ | 0.81[a] | 0.81[a] | 0.81[a] | 0.81[a] |
| $r_{32}$ | 2.10 (3%) | 2.10 (3%) | 2.10 (3%) | 2.09 (2%) |
| $V_H$ (Å$^3$) | 322 (3%) | 317 (3%) | 315 (3%) | 319 (2%) |
| $V_{BW}$( Å$^3$) | 29.86 (0.3%) | 28.12 (0.4%) | 27.97 (0.2%) | 28.36 (0.6%) |
| $a_L$[b] (Å$^2$) | 67.6 (3%) | 69.4 (2%) | 70.2 (2%) | 68.0 (2%) |
| $D_B$[b] (Å) | 37.3 (3%) | 36.3 (2%) | 35.9 (2%) | 37.1 (2%) |
| $D_{pp}$[b] (Å) | 35.4 (3%) | 36.9 (3%) | 37.2 (3%) | 37.1 (3%) |
| $n_W$[b] | $\sim 13$ | $\sim 15$ | $\sim 16$ | $\sim 14$ |
| $\langle\Delta\rangle^{SAXS,b}$ | - | $-0.038$ (60%) | $-0.043$ (60%) | $-0.010$ ($>$ 100%) |
| $N_p$[b] | - | 100 (19%) | 110 (20%) | 150 (13%) |
| $r_p$[b] (Å) | - | 148 (13%) | 138 (14%) | 122 (9%) |

[a] Fixed values

[b] Calculated values

TABLE S8. **DLPC** systems - Summary of fit results for pLUVs with OmpLA **dimers**. Number of iterations $N_I = 100$.

| **Parameter** | LUV mock | PLUV $\hat{x}_p = 1/1200$ | PLUV $\hat{x}_p = 1/300$ |
|---|---|---|---|
| $R_v$ (Å) | 308 (4%) | 370 (8%) | 420 (5%) |
| $Z$ | 19.0 (20%) | 6.8 (20%) | 21 (19%) |
| $x_p^{SAXS}$ | - | 0.00013 (50%) | 0.00046 (20%) |
| $d_{cholCH3}$ (Å) | 1.8 (3%) | 1.79 (3%) | 1.80 (3%) |
| $\delta_{cholCH3}$ (Å) | 3.00 (4%) | 3.00 (3%) | 3.00 (2%) |
| $d_{PCN}$ (Å) | 3.5 (9%) | 3.6 (8%) | 3.9 (8%) |
| $\delta_{PCN}$ (Å) | 2.1 (19%) | 2.5 (16%) | 2.6 (20%) |
| $d_{CG}$ (Å) | 1.00 (11%) | 0.99 (10%) | 1.02 (9%) |
| $\delta_{CG}$ (Å) | 2.20 (9%) | 2.22 (9%) | 2.20 (9%) |
| $2D_C$ (Å) | 20.7 (2%) | 20.6 (1.9%) | 20.8 (2%) |
| $\varepsilon_C$ | 0.08 (9%) | 0.080 (20%) | 0.082 (20%) |
| $\delta_{CH2}$ (Å) | 2.50 (3%) | 2.51 (3%) | 2.52 (2%) |
| $\delta_{CH3}$ (Å) | 1.53 (8%) | 1.53 (7%) | 1.55 (7%) |
| $r_{PCN}$ | 0.259 (3%) | 0.260 (3%) | 0.260 (3%) |
| $r_{CG}$ | 0.473 (3%) | 0.472 (3%) | 0.476 (3%) |
| $r_{32}$ | 1.93 (3%) | 1.93 (3%) | 1.94 (3%) |
| $V_H$ (Å$^3$) | 324 (2%) | 329 (2%) | 322 (3%) |
| $V_{BW}$( Å$^3$) | 30.2 (0.6%) | 29.7 (0.7%) | 30.3 (0.7%) |
| $a_L{}^b$ (Å$^2$) | 64.7 (2%) | 64.6 (2%) | 64.8 (2%) |
| $D_B{}^b$ (Å) | 30.7 (3%) | 30.8 (2%) | 30.7 (2%) |
| $D_{pp}{}^b$ (Å) | 29.1 (3%) | 28.9 (2%) | 29.4 (2%) |
| $n_W{}^b$ | $\sim 8$ | $\sim 8$ | $\sim 9$ |
| $\langle\Delta\rangle^{SAXS,b}$ | - | $-0.004$ (> 100%) | +0.002 (> 100%) |
| $N_p{}^b$ | - | 4 (80%) | 22 (40%) |
| $r_p{}^b$ (Å) | - | 800 (50%) | 370 (20%) |

[b] Calculated values

#### S7. SAXS ANALYSIS - ADDITIONAL PLOTS

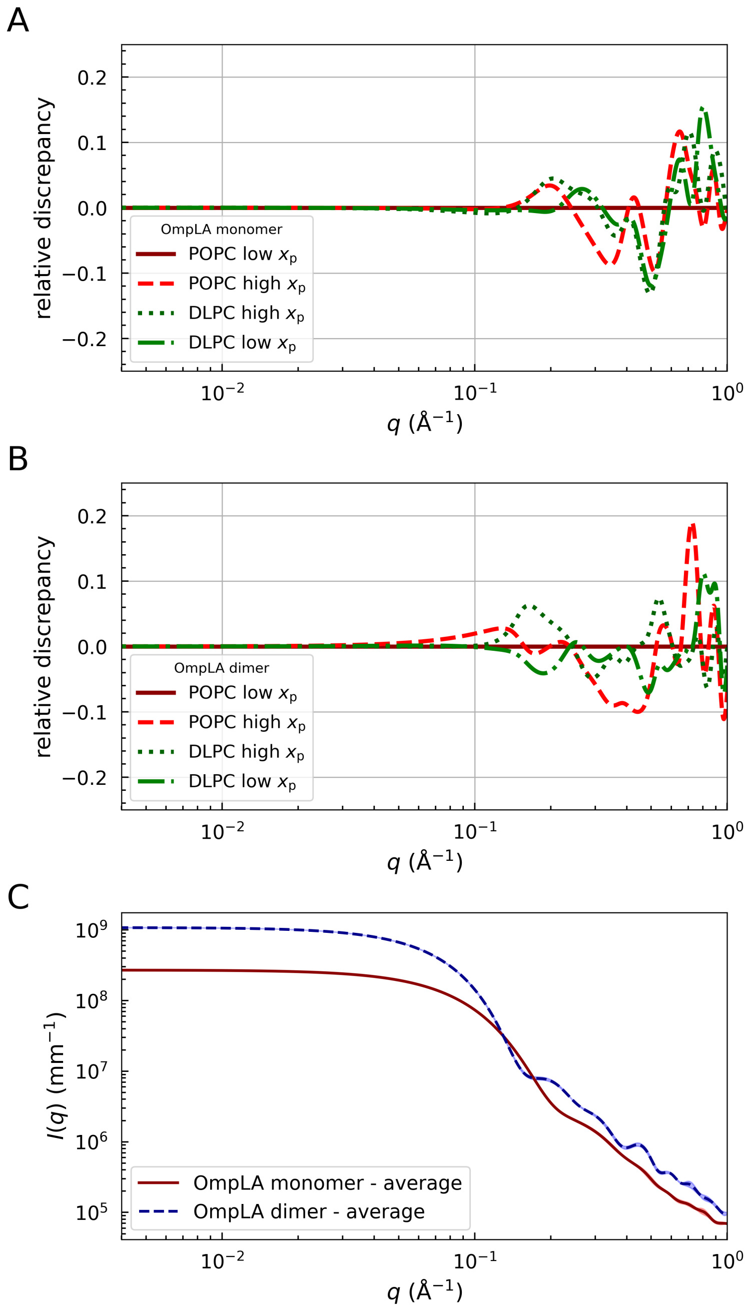


FIG. S3. Comparison of vacuum scattering intensities of OmpLA for different MD configurations from CRYSOL[20]. Panels (A) and (B) show the relative discrepancy between the vacuum scattering intensities of OmpLA monomer and dimer, respectively, in POPC/low-$x_p$ systems and all other configurations, namely POPC/high-$x_p$ (dashed red), DLPC/low-$x_p$ (dotted green), and DLPC/high-$x_p$ (dashed-dotted dark-green). (C) Averaged vacuum scattering intensities of OmpLA monomers (red line) and dimers (dashed blue line).

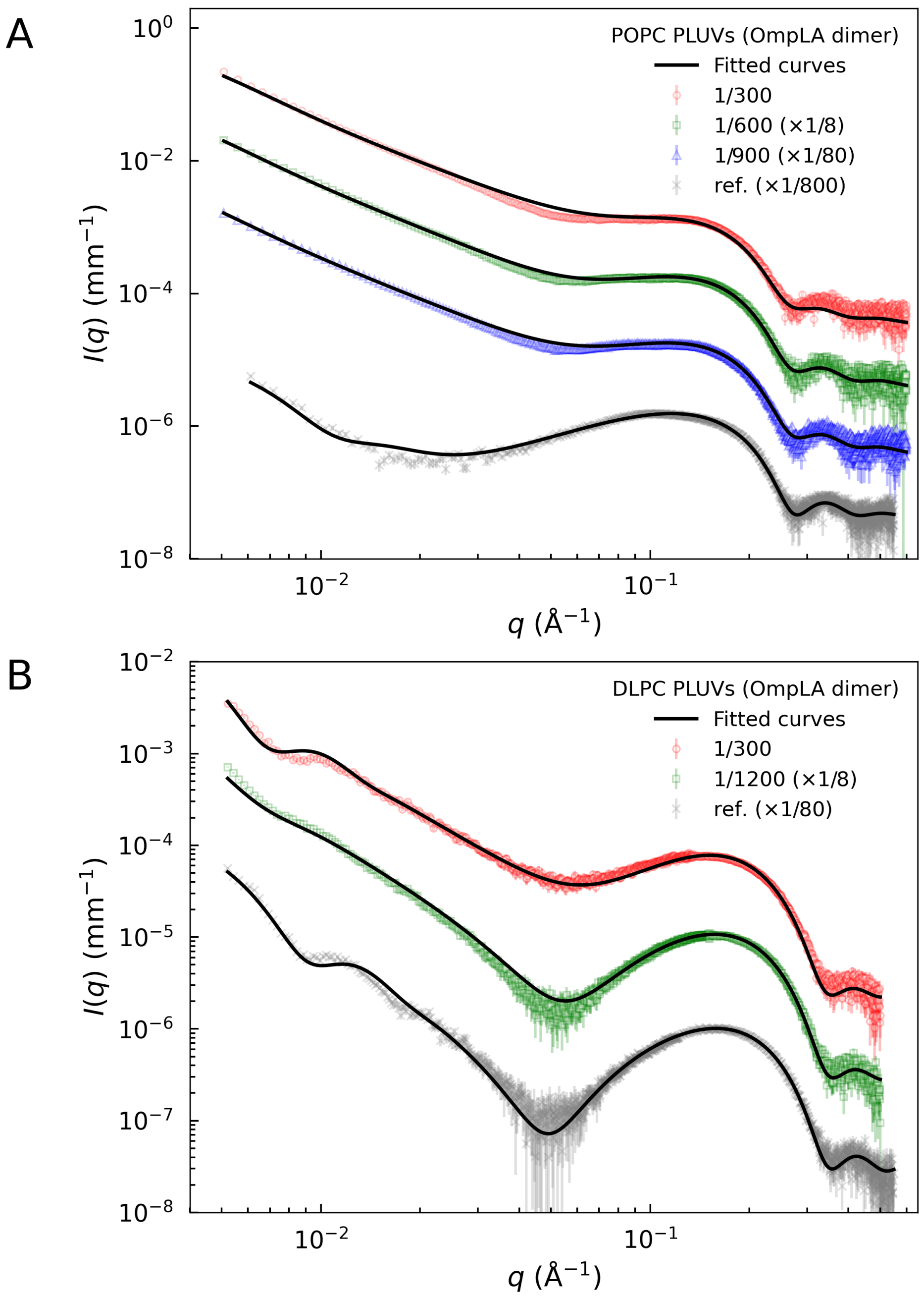


FIG. S4. SAXS data and fitted curves of POPC (A) and DLPC (B) pLUVs at different $\hat{x}_p$ conditions in the case of OmpLA dimers (data and fits were scaled to improve visibility). ‘Mock’ LUV samples serve as references.

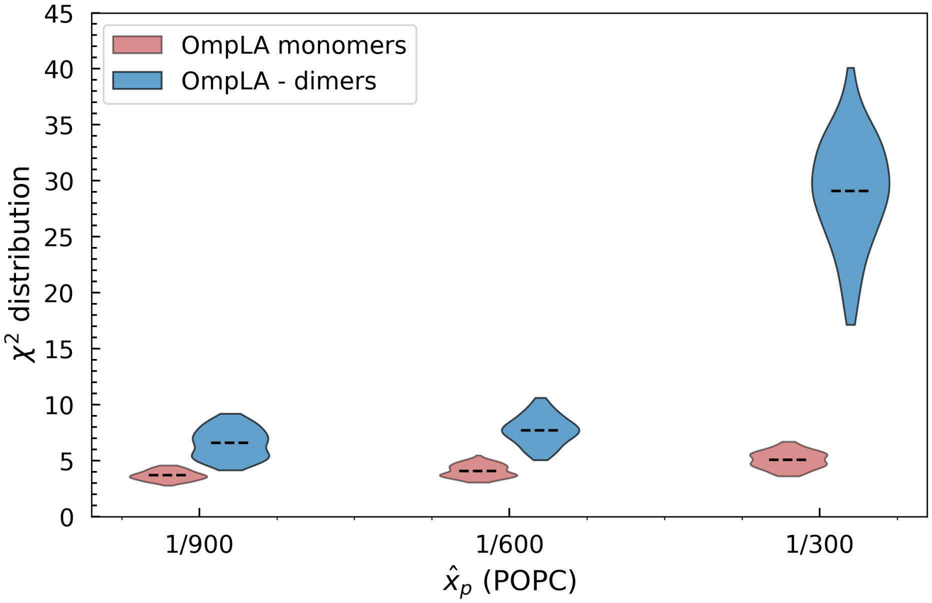


FIG. S5. $\chi^2$ distributions of the POPC pLUV analyses in the case of monomer (red) and dimer (blue) virtual OmpLA models. Dashed lines mark the mean of the distributions.

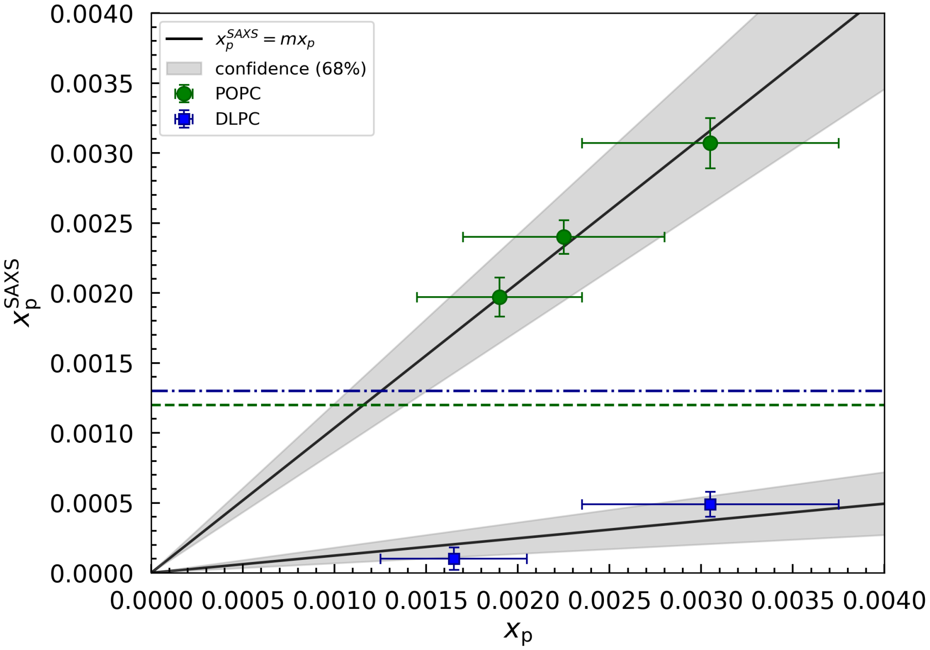


FIG. S6. SAXS analysis results for $x_p^{SAXS}$ as a function of the effective $x_p$ values, assuming a population of OmpLA **dimers**. $x_p$ values and standard deviation were used to define the prior distribution for and $x_p^{SAXS}$. Black line indicate the best fit or $x_p^{SAXS} = mx_p$, while the gray area represents the margin of error at the 68% confidence level. POPC: $m = 1.04 \pm 0.17$, DLPC: $m = 0.12 \pm 0.06$. Dashed and and dotted-dashed lines mark the transition from low to high-$x_p$ regimes for POPC and DLPC, respectively.

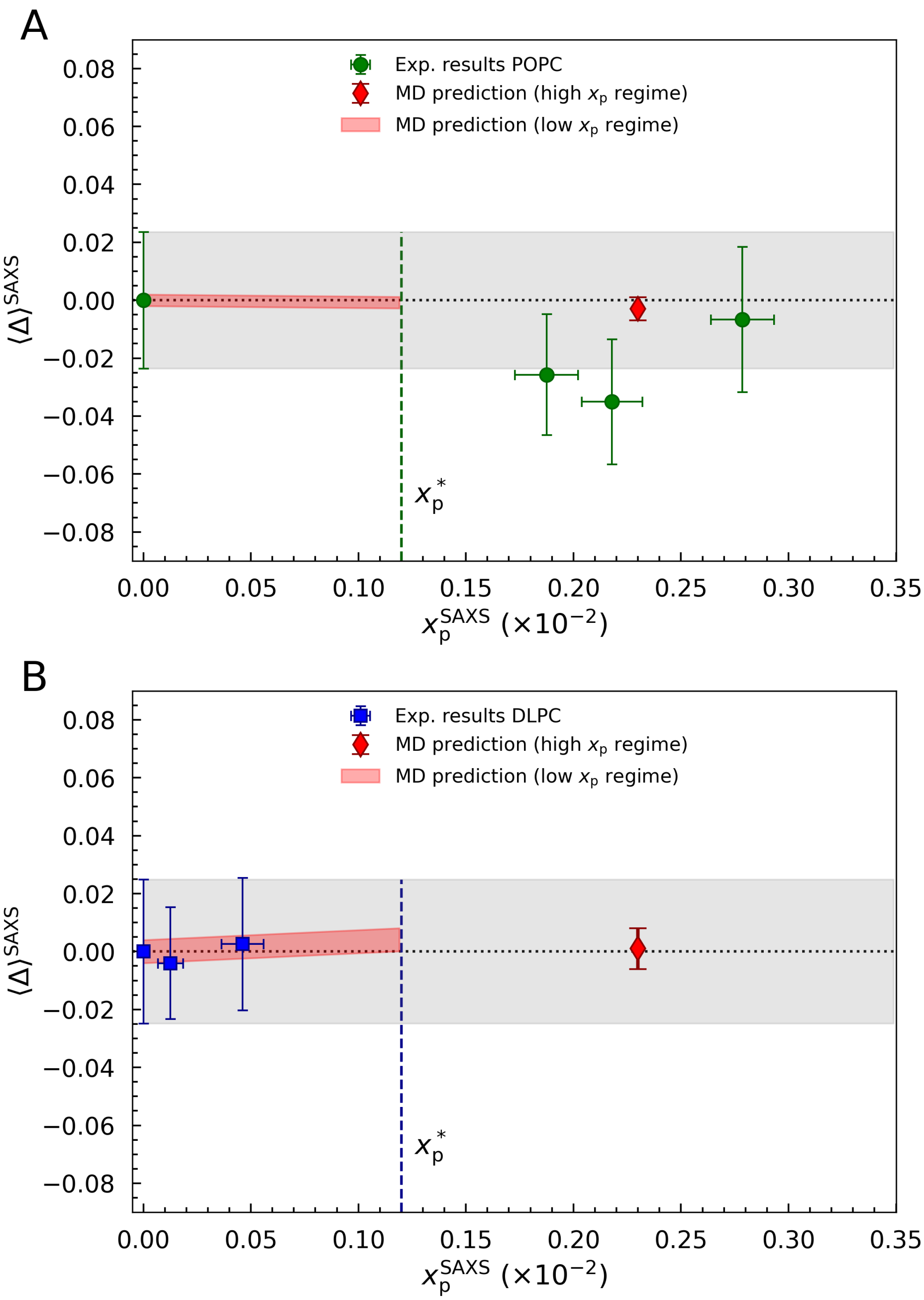


FIG. S7. Average deformation, $\langle\Delta\rangle^{SAXS}$, for POPC (A) and DLPC (B) systems in the case of OmpLA dimers as a function of $x_p^{SAXS}$ results. Dotted lines and gray areas represent reference values for bulk lipids and associated standard deviation. The predicted deformation from MD is shown as red diamonds and red areas for the high-$x_p$ and low-$x_p$ regime, respectively. Vertical dashed lines highlight the transition from low to high-$x_p$ regime.

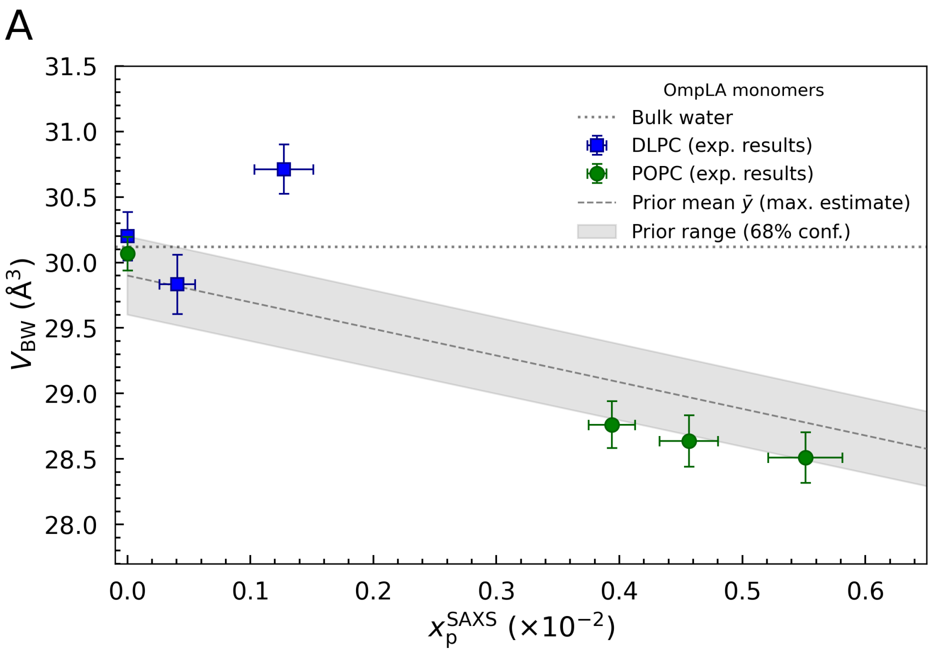


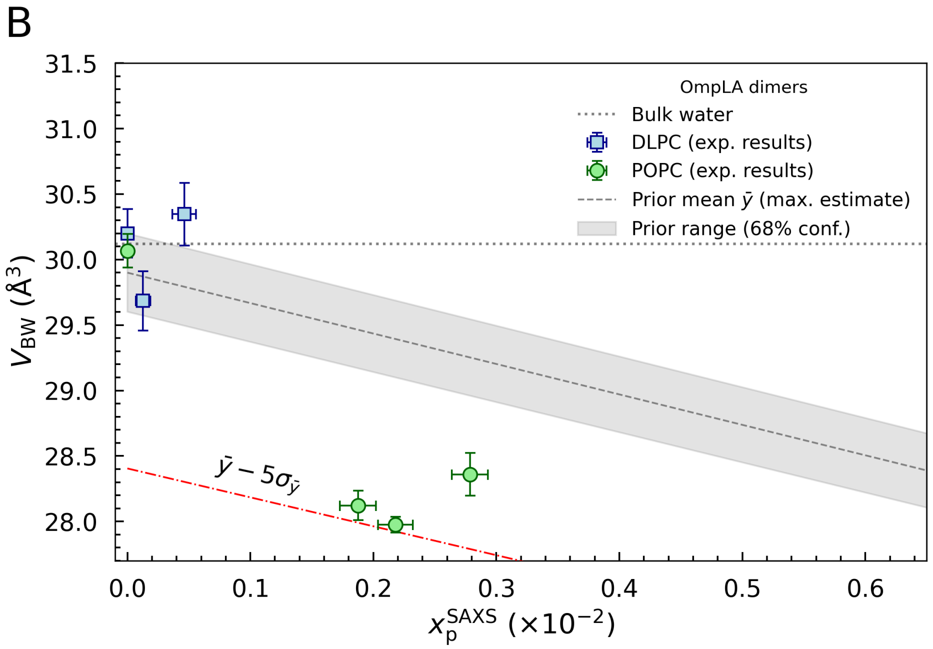


FIG. S8. Mean volume of water molecules in the lipid head-group, $V_{BW}$ for POPC (green circles) and DLPC (blue squares) as a function of $x_p^{SAXS}$ results. Grey areas display enclose the 68% confidence interval of the associate priors mean $\bar{y}$, representing a maximum, possible estimate. The dotted line marks the volume of bulk water molecules. A) Results for OmpLA monomers. B) Results for OmpLA dimers, where the dotted-dashed red line highlights the numerical lower boundary of the Gaussian prior $\mathcal{N}(\bar{y}, \sigma_{\bar{y}}^2)$ at $\bar{y} - 5\sigma_{\bar{y}}$.
